# *P. aeruginosa* disrupts ARP-2/3 mediated apical F-actin organization to induce intestinal deformation in *C. elegans*

**DOI:** 10.1101/2025.10.08.681298

**Authors:** AN Divyashree, Tanushree Sinha, Anup Padmanabhan

## Abstract

Apico-basal polarity is essential for maintaining cytoskeletal organization and regulating transport and secretory functions in the intestinal epithelium. The opportunistic pathogen *Pseudomonas aeruginosa* exploits the interactions between cell polarity and cytoskeletal networks to promote infection. However, most mechanistic insights into these processes have come from *in-vitro* epithelial monolayers, which do not fully capture the complexity of multicellular tissues. Here, we show that heat-labile factor/s secreted by *P. aeruginosa* disrupt apical polarity in *Caenorhabditis elegans* enterocytes through a PI3K-AKT-dependent mechanism. This disruption is accompanied by apical enrichment of PI(3,4,5)P_3_, fragmentation of ARP-2/3 organization, localized disruption of apical F-actin, release of F-actin-rich vesicles into the intestinal lumen, and pronounced deformation of the apical surface. Perturbing CDC-42-ARP-2/3-dependent actin assembly, PI3K-AKT signalling, or the endotube regulators, IFB-2 and IFO-1 suppressed intestinal deformation and significantly extend host survival following *P. aeruginosa* exposure. Notably, these changes occured without detectable loss of epithelial tissue integrity. Together, our findings identify epithelial polarity disruption as a mechanism through which extracellular *P. aeruginosa* remodels the host cytoskeleton and alters tissue mechanics during infection.

**Graphical Abstract:** 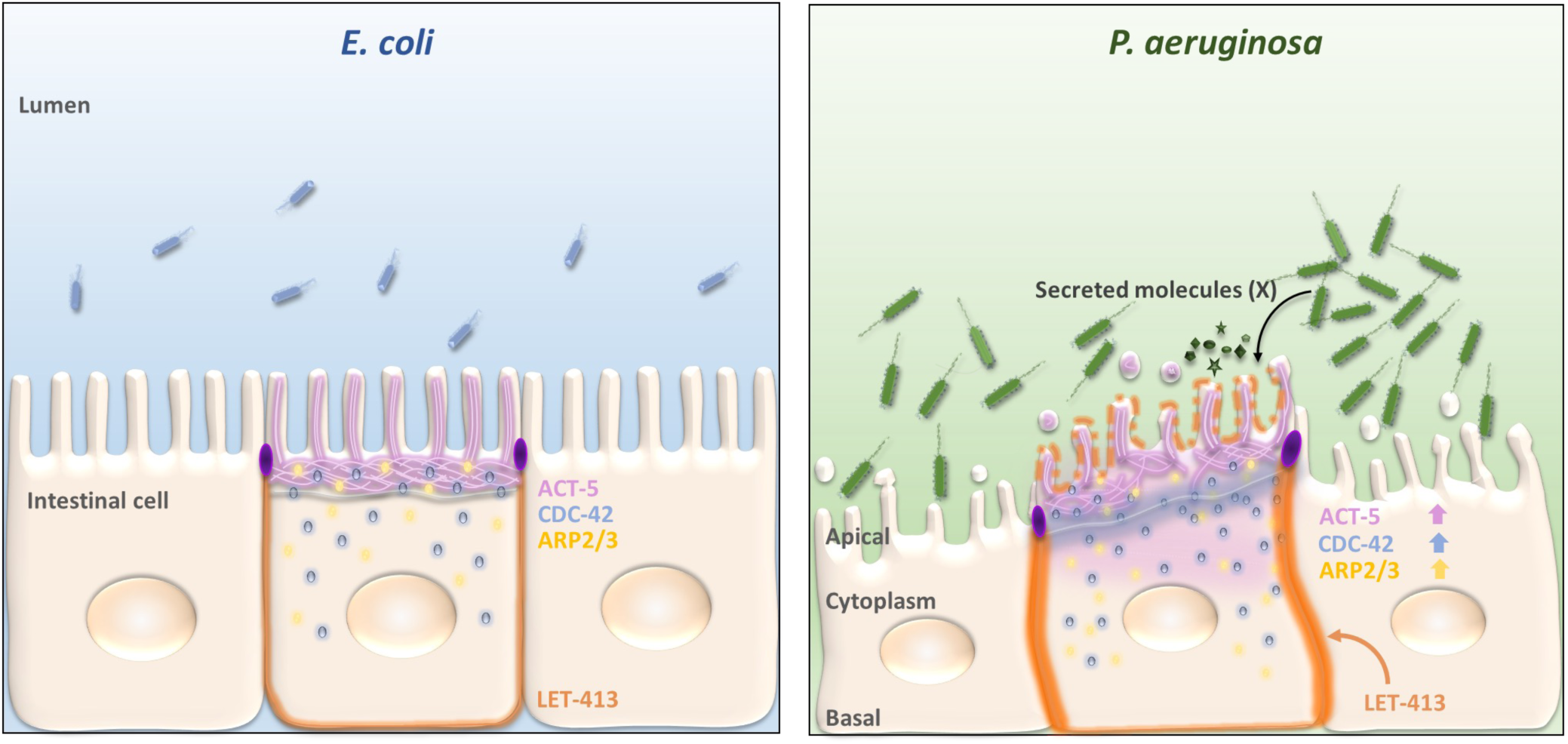

## INTRODUCTION

Constantly exposed to a diverse micro-organisms, the intestinal epithelium acts as a frontline sentinel for immune surveillance-detecting perturbations caused by dysbiosis, distinguishing harmless commensals from pathogens, and mounting context-appropriate immune response. During infection, bacterial pathogens elicit a wide range of responses from intestinal epithelial cells - some orchestrated by the pathogens to promote their survival and dissemination, and others initiated by the host to limit damage and trigger innate immunity. Unlike professional immune cells, enterocytes lack canonical pathogen recognition receptors (PRRs) that detect microbe-associated molecular patterns (MAMPs). Instead, they respond to pathogen effector activity and the disruptions they cause to physiological processes and cellular homeostasis, collectively termed “patterns of pathogenesis”. These include damage associated molecular patterns (DAMPs) triggered by pathogen effectors. Studies in model organisms such as *Drosophila melanogaster* and *Caenorhabditis elegans (C. elegans)* have highlighted the role of effector-triggered, cell autonomous surveillance in epithelial tissue^1–3^.

*C. elegans,* a bacterivorous soil nematode, inhabits microbe-rich environments and encounters a broad spectrum of microorganisms, including pathogens^4,5^. While a protective cuticle^6^ and sophisticated pathogen-avoidance behaviours^7–12^ limits pathogen exposure, the intestinal epithelium remains the primary site of infection due to the worm’s bacterial diet. Ingested bacteria are mechanically ground in the pharynx before entering the lumen, yet some pathogens survive, proliferate, and colonize the intestine. The *C. elegans* intestine is composed of 20 polarized epithelial cells arranged longitudinally, with apical surfaces facing the central lumen and basolateral surface engaged in cell-cell and cell-matrix adhesions^13,14^. The apical surface is enriched in microvilli, providing the primary site for host-pathogen interaction^15^ (Fig. 1A).

**Figure 1:**
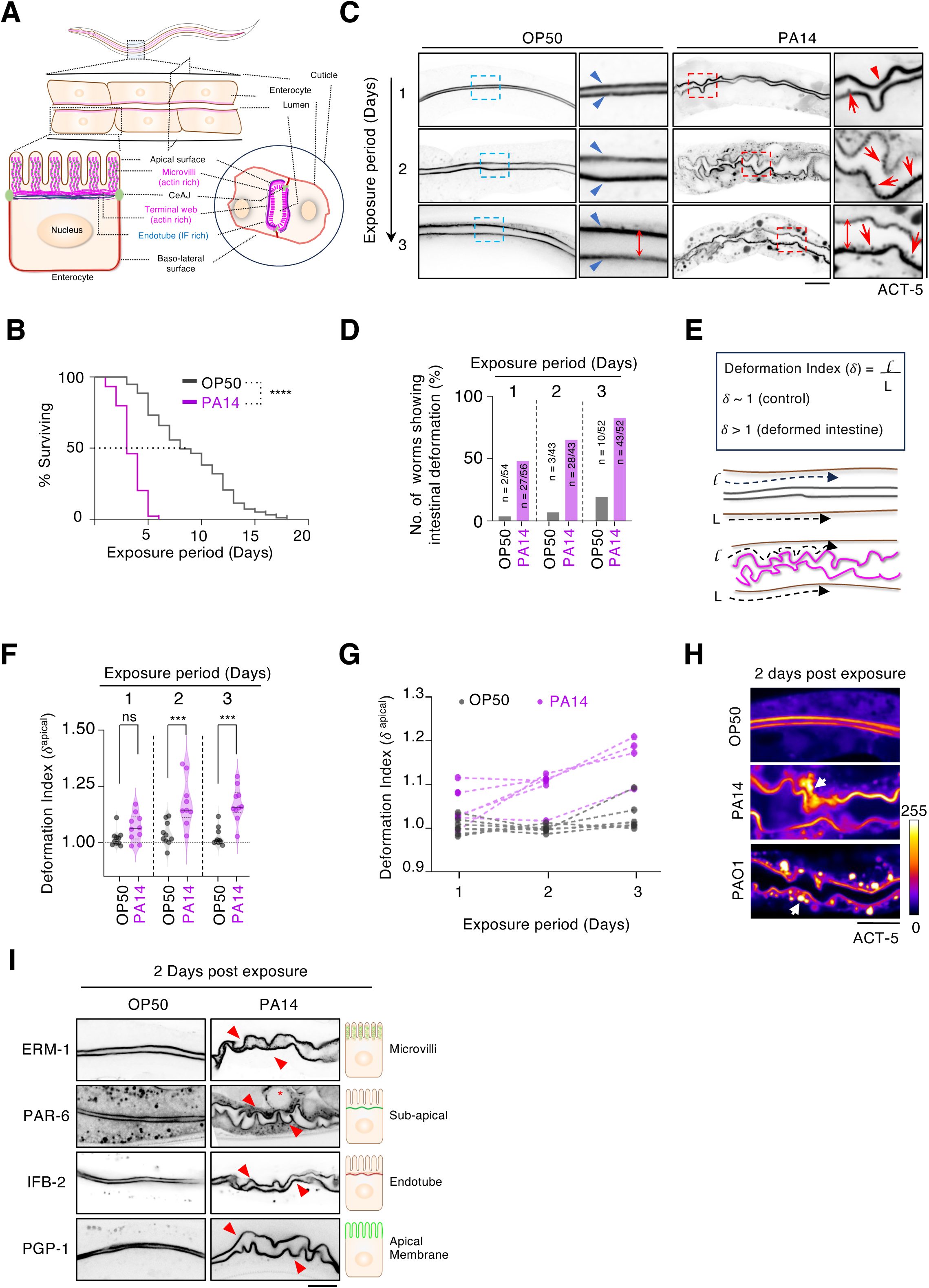
*P. aeruginosa* infection deforms the apical surface of *C. elegans* intestinal epithelium. **(A)** Schematic illustration of *C. elegans* intestinal epithelium highlighting the major subcellular regions of an enterocyte. **(B)** Kaplan-Meier survival curves of N2 worms exposed to *P. aeruginosa* (PA14) or *E. coli*(OP50) at 25⁰ C. (n = 100 worms per condition; N=2) **(C)** Representative medial confocal sections of *C. elegans* intestines expressing mCherry::ACT-5. Although intestinal lumen width increased with age in worms exposed to either *P. aeruginosa* or *E. coli* (red double-headed arrows), apical surface deformation was observed exclusively in *P. aeruginosa*-exposed animals (red filled arrowheads) and was absent in *E. coli*-fed controls (blue ROI; blue filled arrowheads). Actin vesicles budding from the apical surface were frequently observed following *P. aeruginosa* infection (Red ROI; red arrows). **(D)** Quantification showing the percentage of worms exhibiting intestinal deformation following different durations of exposure to *P. aeruginosa*. **(E)** Schematic illustrating the method used to calculate the apical deformation index (*δ*^apical^) corresponding to the conditions shown in (C). ‘*l* ‘ denotes the contour length of the apical membrane, whereas ‘*L*’ denotes the corresponding distance along the worm’s body-axis. **(F)** Quantification of the apical deformation index in worms corresponding to the conditions shown in (C). (n= 10 worms per treatment per experiment; N=3). **(G)** Longitudinal analysis of the apical deformation index in individual worms over three days following exposure to *P. aeruginosa* or *E. coli*. (n=5 worms per treatment) **(H)** Representative medial confocal sections of *C. elegans* intestines expressing mCherry::ACT-5 following exposure to *E. coli* or the *P. aeruginosa* strains (PA14 and PAO1). Apical surface deformation was observed following exposure to both P. *aeruginosa* strains, including the moderately virulent PAO1 isolate. White arrowheads indicate the cytoplasmic accumulation of ACT-5 in infected animals. Images are pseudo colored to highlight differences in fluorescence intensity, with warmer colors indicating higher signal intensity (n≥20). **(I)** Representative medial confocal images of *C. elegans* intestines expressing fluorescently tagged apical and subapical markers upon exposure to *P. aeruginosa* or *E. coli*. ERM-1::eGFP (microvilli associated apical cortex), PAR-6::mNeonGreen (apical PAR polarity complex), IFB-2::mScarlet (intermediate filament network), and PGP-1::GFP (apical plasma membrane) (N=2, n=10). Scale bar is 20 µm. Statistical significance was determined using Mann -Whitney U test. ****p<0.0001, ***p <0.001, and * p <0.05. ‘N’ denotes independent experiments.

The structure and function of the intestinal epithelium depend on the establishment and maintenance of apico-basal polarity. Polarity is maintained by structural and signalling complexes: the Crumbs complex (CRB-1, CRB-3, MAGUK-2) and the PAR complex (comprising PAR-3, PAR-6, PKC-3 kinase and CDC-42 GTPase) localize apically, while the Scribble complex (LET-413, LGL-1, and DLG-1) localize basolaterally. Membrane lipid localizations are similarly polarized, with phosphatidylinositol 4,5-bisphosphate [PI(4,5)P_2_] enriched apically and phosphatidylinositol 3,4,5-trisphosphate [PI(3,4,5)P_3_] restricted to the basolateral surface. This polarity is reinforced by polarity complexes, cytoskeletal organization, membrane trafficking, and junctional assemblies^16–19^. Membrane polarity is also essential for the spatial localization and activation of Rho family GTPases^20^. In particular, CDC-42 activates the ARP-2/3 complex to promote branched actin polymerization, essential for the assembly and maintenance of apical microvilli, enriched in F-actin,, the membrane cytoskeleton linker, ERM-1 and the actin capping factor, EPS-8^21–26^. Beneath the microvillar brush border lies the terminal web, a specialized cytoskeletal structure composed of F-actin, actin-binding proteins, and endotube, a dense intermediate filament network that includes, IFO-1, IFB-2, IFC-2, IFD-2 and IFP-1^27–32^. Together, these cytoskeletal systems provide structural support for the apical surface and maintain epithelial tissue integrity.

During infection many pathogens employ diverse virulence strategies to subvert the host polarity and actin cytoskeletal machinery to alter cellular shape and mechanics^33^. This may facilitate intracellular entry and spread, or extracellular effector-induced inflammation and tissue damage^34,35^. Conversely, epithelial cells may actively reorganize actin as part of their defense response. Thus, plasma membrane polarity-actin cytoskeleton cross talk is vital for epithelial integrity during infection^36,37^.

The opportunistic pathogen, *Pseudomonas aeruginosa* a major cause of nosocomial infections, infects a wide range of hosts, including *C. elegans*^38–40^. In worms, *P. aeruginosa* infection induces diverse stress responses, mitotic quiescence, and apoptosis^41–45^. In the absence of professional immune cells, *C. elegans*, upon detecting infection, activates innate immune signalling such as the TIR-1-p38 MAP kinase and the DAF-2/DAF-16 insulin/IGF-1 pathways^46^. DAF-2 dependent phosphorylation of PI3K/AGE-1 recruits AKT-1/AKT-2 kinases that in turn phosphorylate DAF-16 leading to its cytoplasmic retention. Reduced DAF-2 signalling or disruption of the PI3K-AKT kinase cascade results in translocation of unphosphorylated DAF-16 into nucleus, regulating the gene expression^47,48^. DAF-16 is known to induce expression of antimicrobial genes, detoxifying enzymes, immune genes and bacterial avoidance^49–53^. Avoidance behaviour is also triggered by mechanical cues, such as intestinal distension (“bloating”), mediated by TRPM channels^11,54–56^.

Here, we investigated the mechanism underlying cytoskeletal remodelling and polarity disruption following *P. aeruginosa* infection. We show that heat-labile secreted components by *P. aeruginosa* perturbs CDC-42-ARP-2/3-mediated actin organization in *C. elegans* enterocytes via the PI3K-AKT pathway. This results in the disruption of epithelial polarity and discharges actin rich vesicles from the apical surface leading to deformed intestinal lumen surface and reduced lifespan.

## RESULTS

### *Pseudomonas aeruginosa* infection results in *C. elegans* intestinal deformation

Consistent with previous studies, chronic exposure to *P. aeruginosa* strain PA14 significantly reduces *C. elegans* survival (TD_50_*^PA^*^14^ ∼3 days) compared with *E. coli* (OP50)-fed controls (TD_50_*^OP^*^50^ ∼8 days) at 25⁰ C (Fig. 1B). This lethality has been partly attributed to the colonization of the intestinal lumen by *P. aeruginosa*^40^. To examine the effect of *P. aeruginosa* infection on intestinal cytoskeletal organization, we imaged transgenic *C. elegans* expressing mCherry fused to the N-terminus of ACT-5, the intestine-specific actin isoform that lines the apical surface (Fig. 1A)^15,57^. In *E. coli*-fed worms, ACT-5 appeared as an even, parallel, continuous lining along the intestinal lumen (Fig. 1C; blue arrowhead, OP50). Despite age related intestinal distension^51,56,58^, the ACT-5 signal remained smooth even after 3 days on *E. coli* (Fig.1C; red double-headed arrows in OP50/day-3, and S1A). While exposure to *P. aeruginosa* also resulted in lumen distension (Fig. 1C, red double-headed arrows in PA14 day-3, and Fig S1A), apical surface characteristics were markedly distinct. In contrast to the even surface seen in *E. coli*-fed animals, *P. aeruginosa*-fed worms displayed extensive wrinkles and surface deformations (Fig. 1C; PA14, filled red arrowheads), with vesicle-like structures budding outward into the lumen from the deformed apical membrane (Fig. 1C; PA14, red arrows in enlarged ROI). Compared to only ∼4% (2/54) of *E. coli*-fed worms, ∼48%(27/56) of *P. aeruginosa*-exposed worms showed such deformations after 1-day. By day-3, the incidence rose to ∼83% (43/52) in *P. aeruginosa*-fed worms compared to ∼19% (10/52) on *E. coli* (Fig. 1D).

Magnitude of intestinal deformation depends on the duration *of P. aeruginosa* infection

To quantitatively assess changes in intestinal morphology, we defined a dimensionless “apical deformation index” (*δ*_apical_), calculated as the ratio of apical lumen length to worm body length (Fig. 1E). *E. coli*-fed worms consistently maintained ‘*δ*_apical_^OP50^’∼1, whereas *P. aeruginosa* exposure led to ‘*δ*_apical_^PA14^>1’, that progressively increased with infection duration over two days of exposure and remained elevated thereafter (Fig. 1F). Longitudinal tracking of individual worms further showed that in worms fed with *E. coli*, *δ*_apical_^OP50^ remained ∼1, but increased progressively in worms exposed to *P. aeruginosa* (Fig. 1G). Intestinal deformation was also observed when infection assays were performed at 20⁰ C, ruling out the possibility that the observed phenotype might be a consequence of thermal-stress induced cytoskeletal remodeling (Fig. S1B and S1C). We also observed that a 24-hour exposure to the moderately virulent *P. aeruginosa* strain PAO1 could induce intestinal deformation (Fig. 1H and S1D).

In acute infection assays, 12-hour exposure did not induce significant deformation, whereas 24-hour exposure resulted in higher fraction of worms exhibiting intestinal deformation. Although the 24-hour exposure did not result in statistically significant apical deformation, they persisted even after transfer back to *E.coli* (Fig. S1E). To test whether deformation extended beyond the apical surface, we examined transgenic worms expressing the Scribble homolog, LET-413::mCherry as a marker for the basolateral surface^18,59,17,60^. We detected no significant differences in the basal deformation index (*δ*_basal_), upon exposure to *P. aeruginosa* exposure (Fig. S1F; red arrowhead and S1G). We also excluded the possibility that the intestinal deformation resulted indirectly from increase in oocytes in the germline or uterine expansion due to accumulation of fertilized embryos, by demonstrating similar phenotype in the male intestinal epithelium following *Pseudomonas* exposure (Fig. S1H and S1I). F-actin staining using Phalloidin-AlexaFluor647 (Fig. S1J) and LifeACT::mRuby expression (Fig. S1K) showed visible apical deformation specifically upon *P. aeruginosa* exposure, eliminating the possibility that the infection phenotype might be an artifact of mCherry::ACT-5 transgene expression. We further validated the phenotype using additional apical/sub-apical markers: PGP-1::GFP, ERM-1::GFP, PAR-6::GFP, and IFB-2::GFP, which label the apical membrane, microvilli and subapical positions, respectively (Fig. 1I)^28,61,62^. In all cases, *P. aeruginosa* exposure resulted in clear deformation of the apical surface topology.

The ‘slow killing’ infection assay reported earlier, typically uses nematode growth media (NGM) containing 0.35% peptone^40^. To minimize any confounding effects from elevated peptone concentrations on intestinal epithelial response, we conducted our infections on standard NGM containing 0.25% peptone, optimised for *C. elegans* growth^63^. We observed intestinal deformation in worms infected on 0.35% peptone media indistinguishable from 0.25% peptone (Fig. S1L, S1M and Fig. 1B). Our lifespan assays mirrored previously reported kinetics using 0.35% peptone-containing NGM^40^. Together these findings demonstrate that our infection paradigm faithfully recapitulates chronic *P. aeruginosa* infection, leading to the deformation of the intestinal apical surface.

### CDC-42 mediates *Pseudomonas*-induced intestinal deformation

Members of RHO family GTPases, primarily RHO-1, RAC-1, and CDC-42, are key upstream regulators of F-actin assembly and organization, that coordinate actin dynamics in a spatiotemporally controlled manner^20,64^. In MDCK cells, *P. aeruginosa* infection disrupts RHO signalling through effector proteins that interfere with GTPase-activating proteins (GAPs), leading to the disorganization of actin cytoskeleton^65–69^. To test whether *P. aeruginosa*-induced surface deformation in *C. elegans* enterocytes similarly depends on RHO GTPase-mediated actin re-organization, we individually depleted RHO-1, CED-10 (RAC-1), and CDC-42 in adult worms expressing intestinal mCherry::ACT-5, and subsequently exposed them to *P. aeruginosa* or *E. coli*. Similar to RNAi control animals, both *rho-1(RNAi)* and *ced-10(RNAi)* worms displayed pronounced apical deformation upon *P. aeruginosa* exposure (Fig. 2A, S2A and S2B). In contrast, intestines of *cdc-42(RNAi)* worms exposed to *P. aeruginosa* displayed significantly reduced deformation and appeared remarkably similar to those exposed to *E. coli* (Fig. 2A and 2B). Importantly, this suppression of deformation was not due to impaired pathogen sensing in *cdc-42(RNAi)* worms (Fig. S2C and S2D). Consistent with its role in suppressing intestinal deformation, *cdc-42(RNAi)* worms exhibited significantly extended lifespan (8 days) compared to RNAi controls (4 days) upon *P. aeruginosa* exposure (Fig. 2C).

**Figure 2:**
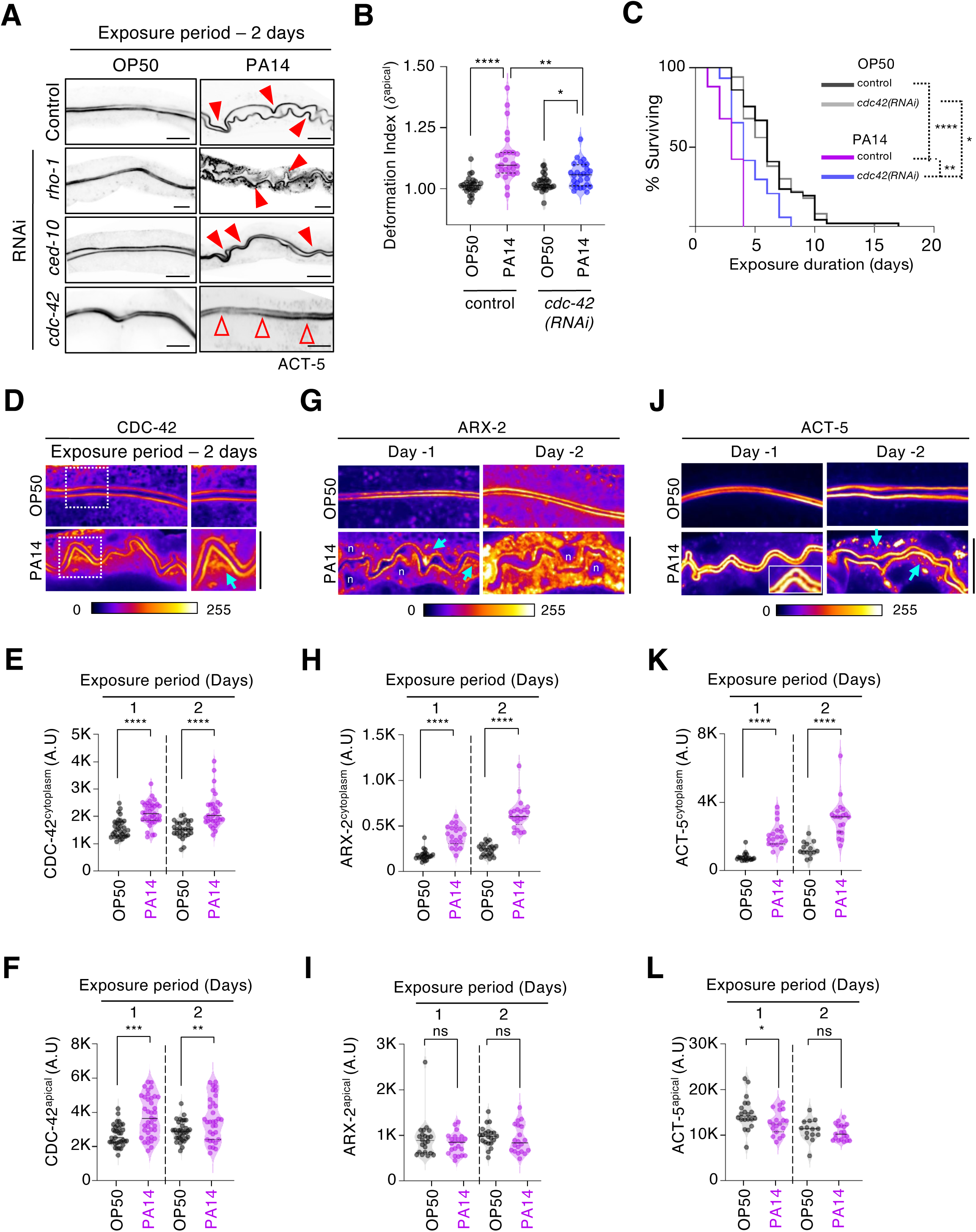
Exposure to *P. aeruginosa* induces cytoplasmic accumulation of CDC-42, ARX-2 and ACT-5 in the C. elegans intestine. **(A)** Representative medial confocal sections of intestines expressing mCherry::ACT-5 following RNAi-mediated depletion of *cdc-42*, *rho-1,* or *ced-10* and subsequent exposure to *P. aeruginosa*. Depletion of CDC-42, but not RHO-1 or CED-10, suppressed *P. aeruginosa* induced intestinal lumen deformation. Red outlined unfilled arrowheads indicate smooth apical surfaces in *cdc-42(RNAi)* worms. **(B)** Quantification of apical deformation index in control (L4440) and *cdc-42(RNAi)* worms following two days of exposure to *P. aeruginosa.* Violin plots show median and interquartile range. (N = 3; n=27) **(C)** Representative Kaplan-Meier survival curves of control (L4440) and *cdc-42(RNAi)* worms following exposure to *P. aeruginosa* or *E. coli*. CDC-42 depletion partially rescued the reduction in lifespan induced by *P. aeruginosa* increasing the mean survival from 4 days in control animals to 8 days in *cdc42(RNAi^)^* animals. (N=2; n= 50 worms per condition per experiment) **(D, G, J)** Representative medial confocal sections of *C. elegans* intestines expressing (D) CDC-42::GFP, (G) ARX-2::TagRFP, or (J) mCherry::ACT-5 following exposure to *E. coli* or *P. aeruginosa*. Images are pseudo-colored to highlight differences in fluorescence intensity, with warmer colors indicating higher signal intensity. Cyan arrows indicate the cytoplasmic accumulation of the respective proteins following *P. aeruginosa* exposure. Where visible, intestinal cell nuclei are labelled ‘n’. **(E and F)** Quantification of (E) cytoplasmic and (F) apical CDC-42::GFP fluorescence intensities in animals exposed to *E. coli* or *P. aeruginosa*. Data are shown as violin plots with mean ±95% C.I. Each dot represents an individual animal. (N=3; n=33 worms). **(H and I)** Quantification of (H) cytoplasmic and (I) apical ARX-2::TagRFP fluorescence intensities in animals exposed to *E. coli* or *P. aeruginosa*. Measurements are presented with mean ±95% C.I. Each dot represents an individual animal. (N=2; n≥21) **(K and L)** Quantification of (K) cytoplasmic and (L) apical mCherry::ACT-5 fluorescence intensities in animals exposed to *E. coli* or *P. aeruginosa*. Measurements are presented with mean ±95% C.I. Each dot represents an individual animal. (N=2; n≥14) Scale bar is 20 µm. Statistical significance was determined using Mann -Whitney U test. ****p<0.0001, ***p <0.001, **p <0.01 and * p <0.05. ‘N’ denotes independent experiments.

### *P. aeruginosa* infection leads to cytoplasmic accumulation of G-actin

The absence of intestinal deformation in *cdc-42(RNAi)* worms prompted us to investigate the intracellular localization of CDC-42 and its downstream effector, the actin nucleating ARP-2/3 complex, during *P. aeruginosa* infection. We used worms co-expressing ARX-2::TagRFP (an essential subunit of the ARP-2/3 complex) at its endogenous locus, and CDC-42::GFP, expressed exogenously under the native *cdc-42* promoter^70,71^. In control animals maintained on *E. coli*, both CDC-42 and ARX-2 localized predominantly to the apical membrane of the enterocytes, with little cytoplasmic accumulation following 2 days of exposure (Fig. 2D and 2G; OP50 panel). Exposure to *P. aeruginosa* led to marked increase in both cytoplasmic and apical CDC-42 levels (Fig. 2D enlarged ROI, 2E, 2F and S2E). Cytoplasmic ARX-2 levels also increased substantially (∼2.6 fold; Fig. 2G and 2H), whereas apical ARX-2 remained largely unchanged relative to *E. coli*-fed controls (Fig. 2I). Intracellular actin mirrored that of ARX-2, with ∼2.5 fold increase in cytoplasmic levels but negligible change in the apical surface (Fig. 2J, 2K and 2L). Interestingly, neither Phalloidin-AlexaFluor647 staining nor LifeACT::mRuby detected cytoplasmic signal, suggesting that the accumulated cytoplasmic pool primarily represents monomeric G-actin (Fig. S1J, S2F and S1K). Together, these findings indicate that *P. aeruginosa* infection promotes CDC-42-dependent cytoplasmic accumulation of G-actin without causing substantial loss of apical F-actin.

### *P. aeruginosa*-induced disruption of cortical actin results in apical surface deformation

It seemed counterintuitive that apical ARX-2 and ACT-5 intensities did not increase despite CDC-42 enrichment following *P. aeruginosa* exposure. To clarify this conundrum, we quantified apical ACT-5 and ARX-2 levels in *cdc-42(RNAi)* worms and found them to be significantly reduced (see OP50 panels in Fig. 3A and S3B, S3C and S3D). Thus, apical actin and ARP-2/3 complex are sensitive to CDC-42-depletion, but not to *P. aeruginosa*-induced CDC-42-enrichment. The absence of deformation in *cdc-42(RNAi)* animals could therefore reflect a direct effect of CDC-42 depletion or indirectly due to reduced ARP-2/3-mediated F-actin polymerization. To distinguish between these possibilities, we depleted ARX-2 or ACT-5 in worms co-expressing CDC-42::GFP and ARX-2::TagRFP. Due to developmental defects associated with *act-5(RNAi)* and *arx-2(RNAi)*, analyses were restricted to day 1 post-infection. Strikingly, *arx-2(RNAi)* and *act-5(RNAi)* worms, like *cdc-42(RNAi)* showed rescue of deformation phenotype (Fig. 3C, 3D, 3E and 3F) following *P. aeruginosa* infection without reduction in cytoplasmic CDC-42 levels (Fig. S3G and S3H). Similarly, pharmacological disruption of ARP-2/3 activity or F-actin polymerization using CK666 or Latrunculin A respectively, prior to exposure to *P. aeruginosa*, resulted in abolishment of the intestinal deformation (Fig. 3G).

**Figure 3:**
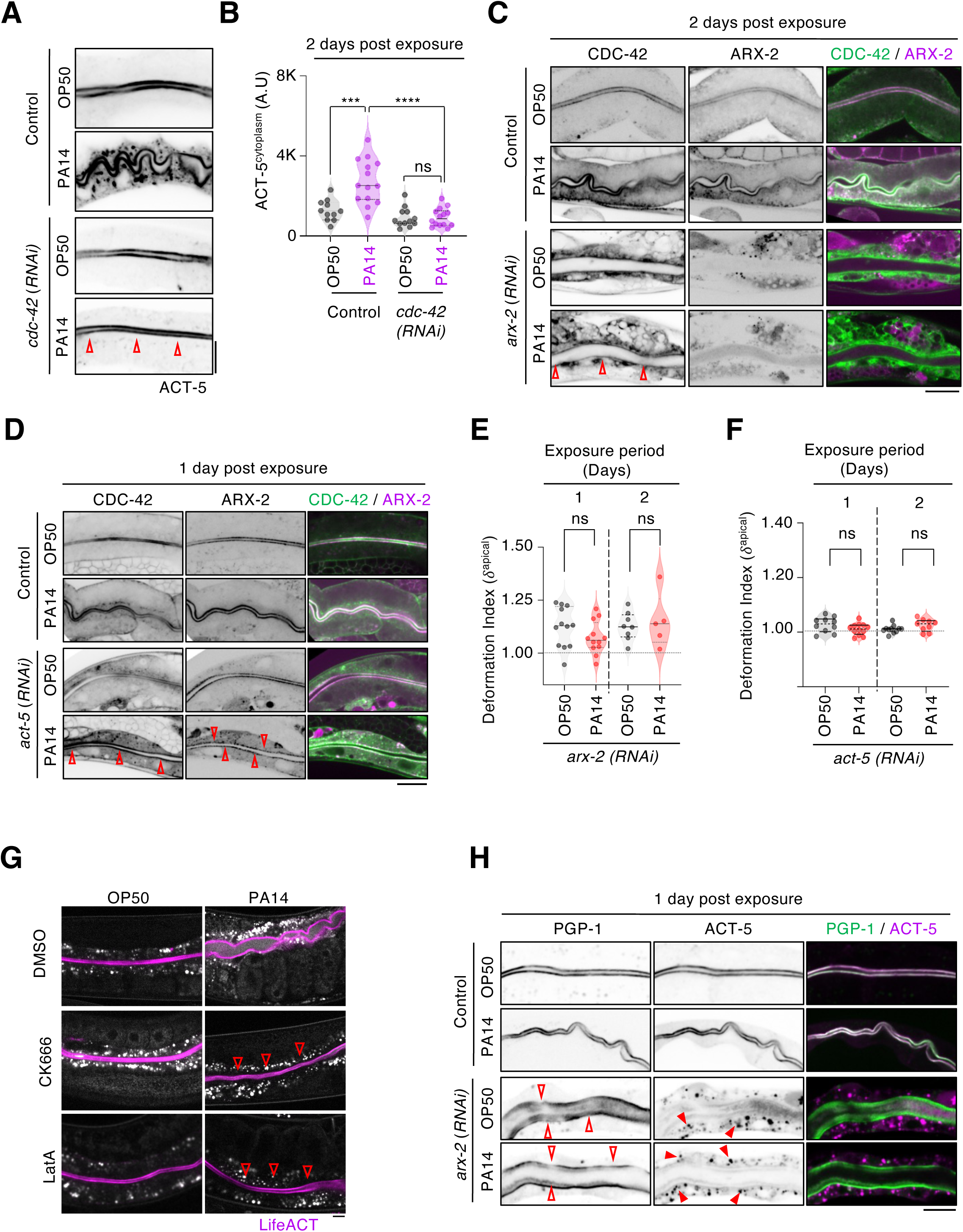
Inhibition of ARP-2/3-dependent actin polymerization suppresses *P. aeruginosa* -induced apical surface deformation. **(A)** Medial confocal sections of Intestinal lumen marked by mCherry::ACT-5 in control (L4440) and *cdc-42(RNAi)* animals following exposure to *P. aeruginosa* or *E. coli*. Red arrowheads indicate suppression of intestinal deformation in *cdc-42(RNAi)* animals. **(B)** Quantification of cytoplasmic ACT-5 fluorescence intensity in the intestinal epithelium of control and *cdc42(RNAi)* worms, two days following exposure to *E. coli* or *P. aeruginosa*. Quantifications are mean ±95% C.I. (N=2, n ≥ 14). **(C)** Confocal images of intestinal epithelium from control and *arx-2(RNAi)* animals co-expressing CDC-42::GFP and ARX-2::tagRFP, following exposure to *E. coli* and *P. aeruginosa*. Images were acquired two days post-exposure. Red outline, zero fill arrows indicate suppression of intestinal deformation in *arx-2(RNAi)* animals. **(D)** Medial confocal planes of intestines co-expressing CDC-42::GFP and ARX-2::TagRFP in control (L4440) and *act-5(RNAi)* animals following exposure to *E. coli* or *P. aeruginosa* Images were acquired one day post-exposure. Red outlined arrows indicate suppression of intestinal deformation in *act-5(RNAi)* animals. **(E)** Violin plots showing quantification of the apical deformation index in *arx-2(RNAi)* animals following exposure to *E. coli* or *P. aeruginosa*. Plot shows the median and interquartile range of each dataset (N=2, n ≥ 11). **(F)** Violin plots showing quantification of the apical deformation index in *act-5(RNAi)* animals following exposure to *E. coli* or *P. aeruginosa*. Plot shows the median and interquartile range of each dataset (n ≥ 10). **(G)** Medial confocal sections of the intestinal lumen labelled with LifeAct:mRuby in worms treated with CK-666 or latrunculin A following exposure to *E. coli* or *P. aeruginosa*. **(H)** Medial confocal sections of intestines co-expressing PGP-1::GFP and mCherry::ACT-5 in control (L4440) and *arx-2(RNAi)* animals following exposure to *E. coli* and *P. aeruginosa*. Images were acquired one day post-exposure. Scale bar is 20 µm. Statistical significance was determined using Mann -Whitney U test. ****p<0.0001, ***p<0.001, and * p <0.05. ‘N’ denotes independent experiments.

To further investigate the role of ARP-2/3-dependent actin polymerization in *P. aeruginosa*-induced intestinal deformation, we examined worms co-expressing mCherry::ACT-5 and the apical membrane maker, PGP-1::GFP. In control worms fed with *E. coli*, ACT-5 and PGP-1 uniformly co-localized along the apical surface (Fig. 3H; control panel). In contrast, *arx-2(RNAi)* intestines lacked organized apical actin which instead accumulated as cytoplasmic aggregates (Fig. 3H; *arx-2(RNAi)* red filled arrowheads). This observation ruled out cytoplasmic actin accumulation as a possible cause of *P. aeruginosa*-induced deformation. PGP-1 displayed diffused apical localization in *arx-2(RNAi)* intestines (Fig. 3H; *arx-2(RNAi)* red hollow arrowheads), highlighting the essential role of ARP-2/3 polymerized actin in maintaining apical membrane organization. Importantly, CDC-42 and ARX-2 maintained apical localization even in the absence of actin (Fig. 3D, S3I and S3K), confirming their upstream role in the actin polymerization pathway. Collectively, these results demonstrate that suppression of apical deformation in *cdc-42(RNAi)* and *arx-2(RNAi)* intestines following *P. aeruginosa* exposure was due to impaired ARP-2/3 mediated actin polymerization.

### Fragmented ARP-2/3 clustering and actin disorganization drive release of F-actin-rich apical membrane vesicles into the intestinal lumen following *P. aeruginosa* exposure

Although apical intensities of ARX-2 and ACT-5 were comparable between worms exposed to *P. aeruginosa* and *E. coli* (Fig. 2I and 2L), their spatial organization at the apical differed markedly. In *E. coli*-fed worms, ARX-2 was evenly distributed along the apical surface (Fig. 4A; OP50, cyan arrowheads), consistent with its established role in maintaining contiguous actin polymerization and brush border integrity^15,72,73^. In contrast, *P. aeruginosa*-infected animals exhibited ARX-2-rich clusters, interspersed with ARX-2-depleted regions (Fig. 4A; PA14, white arrows). We sought to correlate this fragmented ARP-2/3 localization with the cortical actin distribution using worms co-expressing mCherry::ACT-5 and ARX-2::GFP. Medial confocal sections revealed prominent actin-rich protrusions emerging from ARX-2 depleted region at the apical surface and extending into the intestinal lumen (Fig 4B, enlarged ROI; PA14 panel, yellow arrowheads,- actin protrusions, cyan arrows-fragmented ARP-2/3 and Fig. 1C; red arrows in enlarged ROI). By day-2, these protrusions pinched off as extracellular vesicles into the lumen(Fig. 1C; days 2 & 3, PA14 panel-red arrows in enlarged ROI). This was in contrast to the uniform and undisturbed actin in *E-coli*-fed worms (OP50 panel in Fig. 1C and Fig. 4B). Lattice light sheet microscopy of the intestinal lumen surface of animals expressing mCherry::ACT-5 exposed to *P. aeruginosa* expressing GFP revealed extracellular pinching of actin rich vesicles from the apical surface into the lumen (Fig. 4C; enlarged ROI, yellow arrows). These vesicles were also labelled by LifeACT::mRuby, confirming the presence of filamentous F-actin within these structures (Fig. 4D).

**Figure 4:**
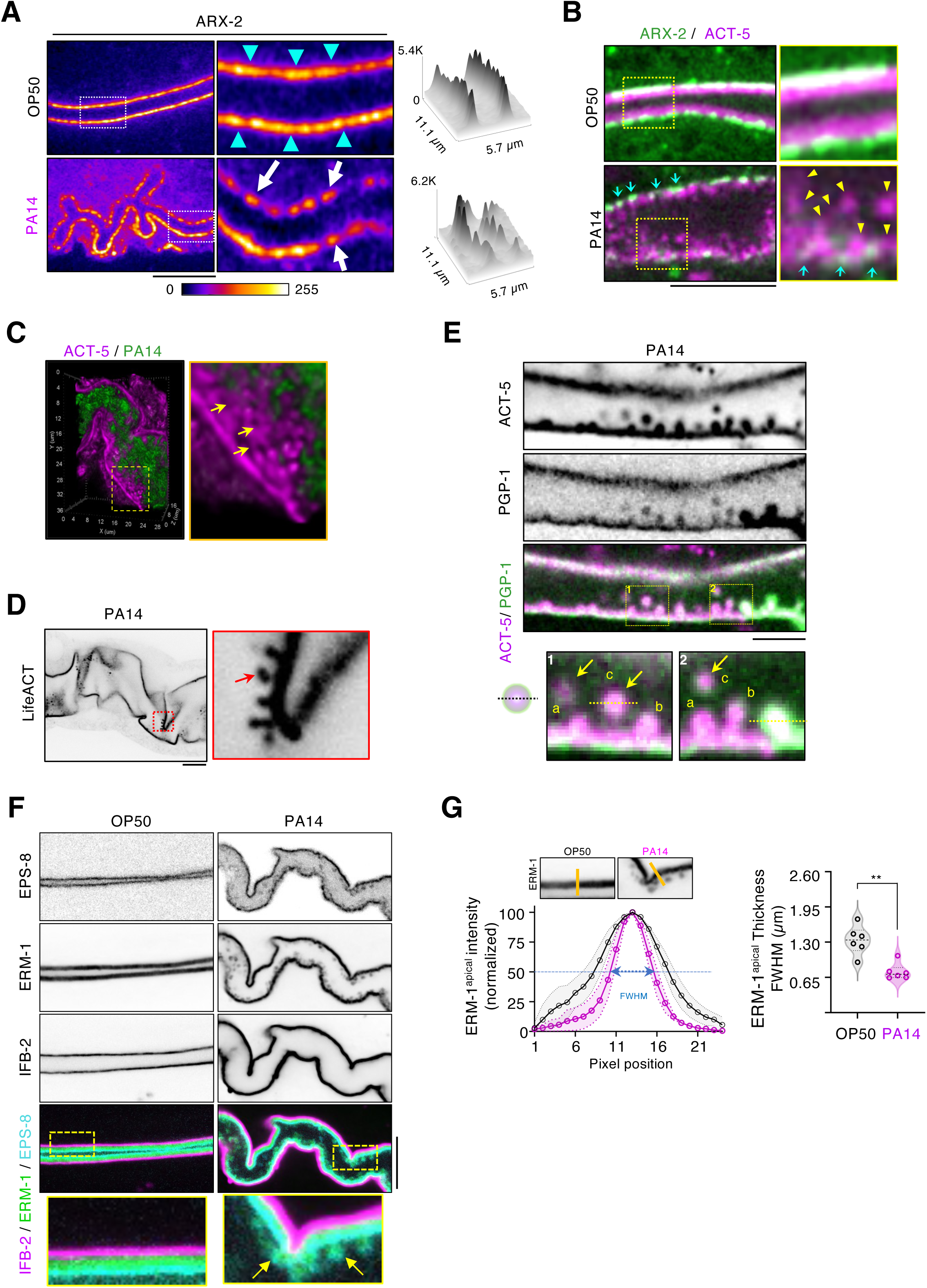
Fragmentation of ARP-2/3 organization disrupts apical actin architecture and promotes extracellular vesicle release from the intestinal surface. **(A)** Intensity-rendered medial confocal images of *C. elegans* intestines expressing ARX-2::TagRFP two days following exposure to *E. coli* or *P. aeruginosa*. In *E. coli*-fed animals, ARX-2 exhibited contiguous apical localization (cyan arrowheads), whereas *P. aeruginosa* exposure resulted in fragmented ARX-2 localization (white arrows). Surface intensity plots of the enlarged regions of interest (ROIs) highlight the discontinuous distribution of ARX-2 following infection. Scale bar −10 µm. **(B)** Confocal images of intestines co-expressing ARX-2::GFP and mCherry::ACT-5 showing actin-rich structures (yellow arrowheads) extending into the intestinal lumen from regions depleted of ARX-2 (cyan arrows), suggesting local disruption of ARP-2/3-dependent actin organization. Scale bar −10 µm. **(C)** Lattice light sheet images of worms expressing intestinal mCherry::ACT5 and exposed to PA14-GFP or OP50-GFP for two days. Acquired Images are represented as maximum intensity projection rendered in volume mode. Yellow arrows in the enlarged ROI indicate vesicles emerging from the apical surface into the intestinal lumen. **(D)** Confocal image of intestines expressing LifeACT::mRuby following *P. aeruginosa* exposure. Extracellular vesicles present within the intestinal lumen are F-actin-rich (red arrow). Scale bar - 10 µm. **(E)** Membrane-bound, actin rich vesicles released from the intestinal apical surface following two days of *P. aeruginosa* exposure. Yellow arrows in the ROI indicate vesicles associated with the apical membrane at various stages of budding – (a) initial membrane protrusion, (b) bud formation, and (c) released vesicle. A fluorescence intensity line scan across the extracellular vesicle demonstrates actin-rich core labelled by mCherry::ACT-5 (magenta) surrounded by the apical plasma membrane marked PGP-1::GFP (green). See Fig. S4A for quantification. Scale bar - 5 µm. **(F)** Medial confocal sections of the *C. elegans* intestine co-expressing ERM-1::mNeonGreen, IFB-2::mScarlet, and EPS-8::BFP-2 following two days of *Pseudomonas* exposure. Yellow arrows in the enlarged ROI indicate the ERM-1-positive vesicles budding from the apical surface of infected intestines. Scale bar - 10 µm. **(G)** Line scan-intensity plot of ERM-1::mNeonGreen the intestines of *C. elegans* exposed to *P. aeruginosa* or *E. coli* for two days. Plot of the full width at half maximum (FWHM) demonstrates reduction in apical ERM-1 width following *P. aeruginosa* infection. Statistical significance was determined using Mann -Whitney U test. **p <0.01

To determine whether these extracellular F-actin structures were membrane-enclosed, we examined worms co-expressing mCherry::ACT-5 and PGP-1::GFP. Medial confocal images on day-2 post-infection revealed multiple stages of vesicle biogenesis: initial protrusion (Fig. 4E; ROIs 1a & 2a), membrane pinching during vesicle budding (1b & 2b), and free, detached vesicles within the lumen (yellow arrows, 1c & 2c). Intensity line-scan analysis further confirmed that these vesicles, measuring ∼ 0.62± 0.01µm in diameter, contained a central F-actin core surrounded by PGP-1::GFP-positive apical membrane (Fig. S4A and S4B). Previous studies have shown that phosphorylation of ACT-5 by the JNK-family kinase KGB-1 promotes cytoplasmic redistribution of ACT-5 in ageing worms, leading to intestinal dysbiosis^58^. We therefore investigated whether KGB-1-dependent actin remodelling might contribute to the vesicle release and deformation phenotype during *P. aeruginosa* infection. However, depletion of KGB-1 failed to suppress *P. aeruginosa*-induced intestinal deformation (Fig. S4C), suggesting that cytoskeletal remodelling described here is independent of KGB-1-mediated phosphorylation of ACT-5.

### Extracellular vesicles released during *P. aeruginosa* infection originate from apical microvillar surface

Brush border microvilli define the apical surface of enterocytes in many animals, including *C. elegans*^14^. These highly ordered membrane protrusions exhibit uniform dimensions and are enriched in F-actin as well as various actin-binding and regulatory proteins^74^. We therefore asked if the extracellular F-actin containing vesicles released into the lumen following *Pseudomonas* exposure originated from microvillar protrusions. To address this, we used *C. elegans* co-expressing fluorescent protein fusions to the F-actin barbed-end capping protein EPS-8, membrane-cytoskeletal linker ERM-1, and the intermediate filament protein, IFB-2^72^. EPS-8 and ERM-1 localize to the tips and shafts of the microvilli, respectively, whereas IFB-2 localizes to the endotube at the base of microvilli (Fig. 4F; OP50 panel and Fig. 1A)^24,26,61,62,72^. As a proxy for microvillar length, we measured the full width at half maximum (FWHM) of ERM-1 fluorescence, which was ∼1.34μm in *E. coli*-fed worms (Fig. 4G; OP50-grey curve). Following exposure to *Pseudomonas*, both ERM-1 and EPS-8 were detected within extracellular vesicles released into the lumen, whereas IFB-2 (marking intermediate filaments) was absent (Fig. 4F; PA14 panel). Concomitantly, microvillar length was reduced by 50%, with ERM-1 FWHM decreasing to ∼ 0.71μm (Fig. 4G; PA14-magenta). The observation that the extracellular vesicles contained microvillar components, including F-actin, ERM-1 and EPS-8, but lacked terminal web-associated proteins such as CDC-42, ARP-2/3, and IFB-2, suggests that *P. aeruginosa* infection induces progressive disassembly of microvillar material as extracellular vesicles into the intestinal lumen.

### Depletion of IFO-1 and IFB-2 promotes tissue-wide release of apical CDC-42-ARP2/3-F-actin containing extracellular vesicles that are distinct from *P. aeruginosa*-induced microvillar vesicles

Intermediate filaments are closely associated with the apical F-actin network and contribute to the structural integrity of the intestinal cortex^28,29^. Consistent with this, the endotube marked by IFB-2 exhibited pronounced deformation following *P. aeruginosa* exposure (Fig. 1I and 5A). Moreover, both apical and cytoplasmic IFB-2 levels were significantly elevated in infected animals (Fig. 5A and 5B). This infection-induced increased IFB-2 was abolished upon ARX-2 depletion. (Fig. S5A, and S5B). *arx-2(RNAi)* animals, whose intestinal deformation was significantly diminished upon *P. aeruginosa* exposure, also displayed control-like IFB-2 (Fig. 5C, S5A, and S5B). To investigate the contribution of intermediate filaments to intestinal deformation, we quantified ‘*δ*^apical^’ in worms expressing mCherry::ACT-5 following RNAi-mediated depletion of *ifo-1*, *ifb-2*, *ifc-2*, *ifd-1*, or *ifp-1* and subsequent exposure to either *E. coli* or *P. aeruginosa*. Similar to control, *ifc-2(RNAi)*, *ifd-1(RNAi)*, and *ifp-1(RNAi)* animals displayed extensive intestinal deformation upon *P. aeruginosa* infection. In contrast, depletion of either *ifo-1* or *ifb-2* markedly suppressed pathogen-induced deformation Fig. 5D and 5E). We ruled out the possibility that the reduced deformation was due to the effects of RNAi-mediated intermediate filament depletion on body size (Fig. S5F). Despite reduced intestinal deformation, extracellular F-actin-positive vesicles were still detected in *ifo-1(RNAi)* and *ifb-2(RNAi)* animals following *P. aeruginosa* exposure (Fig. 5D; ROI inset in *ifo-1(RNAi)* and *ifb-2(RNAi)*-PA14). Intriguingly, we detected F-actin containing extracellular vesicles within the intestinal lumen of *ifb-2(RNAi)* animals exposed to *E. coli* as well (Fig. 5D; ROI inset in *ifb-2(RNAi)*-OP50).

**Figure 5:**
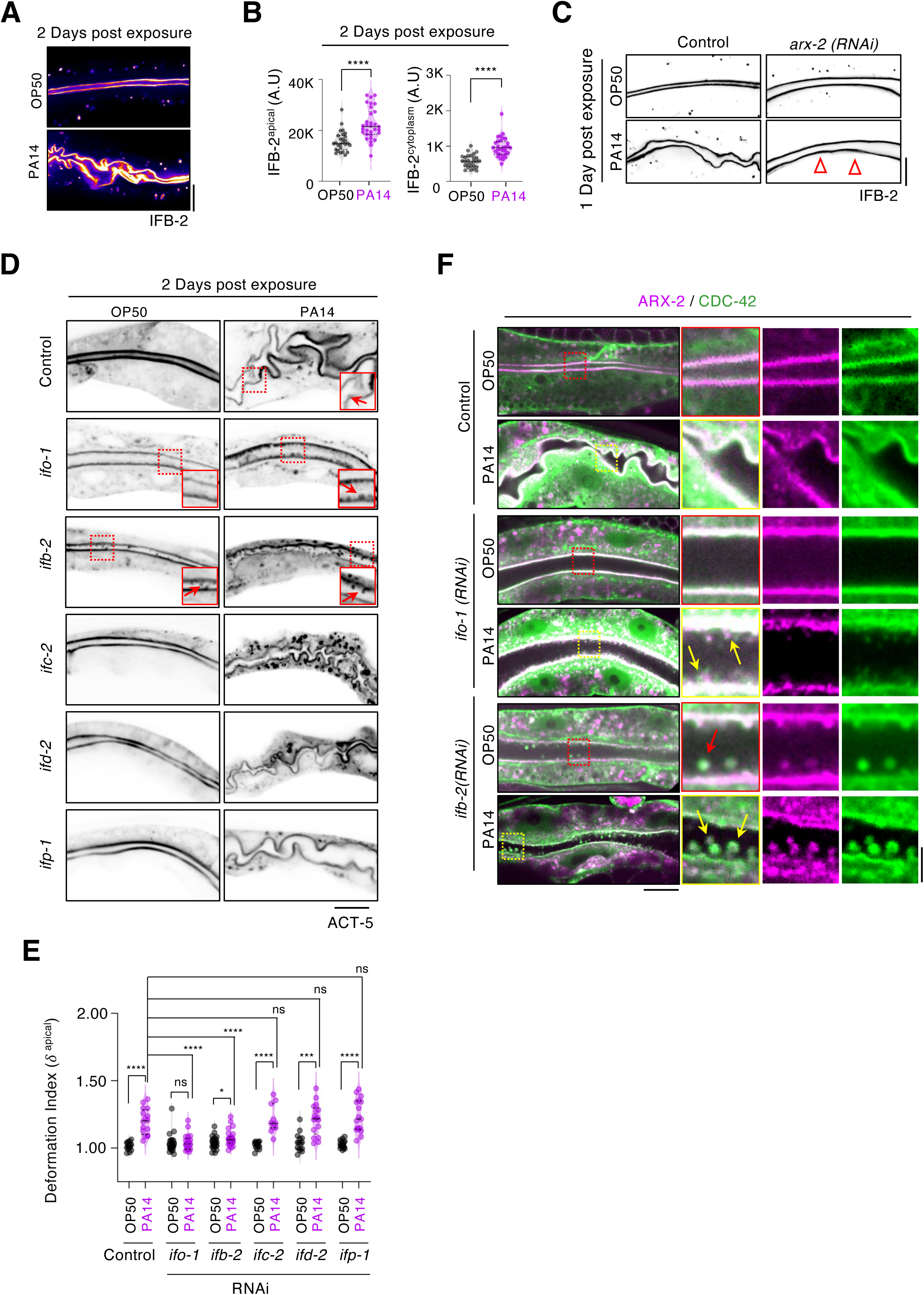
Intermediate filaments regulate intestinal deformation and extracellular vesicle composition. **(A)** Medial confocal sections of the *C. elegans* intestines expressing IFB-2::mScarlet following two days of exposure to *E. coli* or *P. aeruginosa*. Images are pseudo colored to highlight differences in fluorescence intensity with warmer colors indicating higher signal intensity. **(B)** Violin plots showing quantification of apical and cytoplasmic IFB-2::mScarlet fluorescence intensity following two days exposure to *E. coli* or *P. aeruginosa*. Measurements are presented as mean ±95% C.I. Each dot represents an individual animal (N=2; n≥27). **(C)** Medial confocal image of intestinal lumen expressing IFB-2::mScarlet in control (L4440) and *arx-2(RNAi)* animals following exposure to *E. coli* or *P. aeruginosa.* Images were acquired one day post-exposure. Red outline, zero fill arrowheads indicate suppression of intestinal deformation in *arx-2(RNAi)* animals. **(D)** Medial confocal sections of intestines expressing mCherry::ACT-5 following RNAi-mediated depletion of intestinal intermediate filament genes. Depletion of *ifo-1 and ifb-2*, but not *ifc-2, ifd-2 or ifp-1* suppressed *P. aeruginosa*-induced intestinal deformation. Red arrows in insets highlight ACT-5-positive lumenal vesicles. **(E)** Plots showing quantification of apical deformation index in control(L4440) and animals depleted of intestinal intermediate filaments proteins. Depletion of IFO-1 and IFB-2 reduced intestinal deformation, whereas depletion of IFC-2, IFD-2, and IFP-1 has negligible effect. Plots show the median and interquartile range of each data set. Each dot represents an individual worm (N= 2; n≥21 for *ifo-1* and *ifb-2* ; n≥11 for *ifc-2, ifd-2*, and *ifp-1*). **(F)** Medial confocal sections of *C. elegans* intestines co-expressing CDC-42::GFP and ARX-2::tagRFP following two days of exposure to *E. coli* or *P. aeruginosa*. Yellow arrow in the enlarged ROI indicate extracellular vesicles positive for CDC-42::GFP and ARX-2::tagRFP budding from the apical surface into the intestinal lumen. Scale bar in 20 µm. Statistical significance was determined using Mann -Whitney U test. ****p<0.0001, ***p <0.001, **p <0.01, and * p <0.05. ‘N’ denotes independent experiments.

We next investigated the converse scenario, where we depleted IFO-1 and IFB-2 in worms co-expressing fluorescently tagged ARX-2 and CDC-42, before infecting them with *P. aeruginosa*. As seen earlier, exposure to *P. aeruginosa* did not cause intestinal deformation in *ifo-1(RNAi)* and *ifb-2(RNAi)* animals. Correspondingly, *P. aeruginosa*-induced cytoplasmic accumulation of ARX-2 was suppressed in *ifo-1(RNAi)* animals, whereas apical ARX-2 levels remained largely unchanged (Fig. S5C and S5D). Despite the absence of intestinal deformation, we observed extracellular vesicles in the lumen of these worms (Fig. 5F; yellow arrows in enlarged ROIs), positive for ARX-2 and CDC-42. This is in contrast to extracellular vesicles released following *P. aeruginosa* infection in control animals, which were F-actin containing, but devoid of CDC-42 and ARP-2/3 (Fig. 4B, and 5D). Thus, disruption of the intermediate filament-rich endotube triggers tissue-wide shedding of apical CDC-42–ARP-2/3–F-actin-containing membrane domains, producing a class of extracellular vesicles that is distinct in origin and composition from microvillar vesicles induced by *P. aeruginosa* infection. Together, these findings highlight the essential function of intermediate filament architecture in the organization of apical surface in *C. elegans* intestine.

### *P. aeruginosa* infection disrupts apical polarity of the enterocyte epithelium

In MDCK cells, enhanced PI(3,4,5)P_3_ at the apical surface following *P. aeruginosa* attachment results in the loss of apical membrane identity^75–77^. Fragmented ARP-2/3 clustering and vesicle shedding following *P. aeruginosa* exposure suggested loss of apical membrane identity. Consistent with these observations, intestines from worms exposed to *P. aeruginosa,* exhibited elevated PI(3,4,5)P_3_ levels compared to *E-coli*-fed worms (Fig. 6A). Animals treated with wortmannin prior to *P. aeruginosa* exposure prevented the infection-induced increase in the intestinal PI(3,4,5)P_3_ levels (Fig. 6A and S6A).

**Figure 6:**
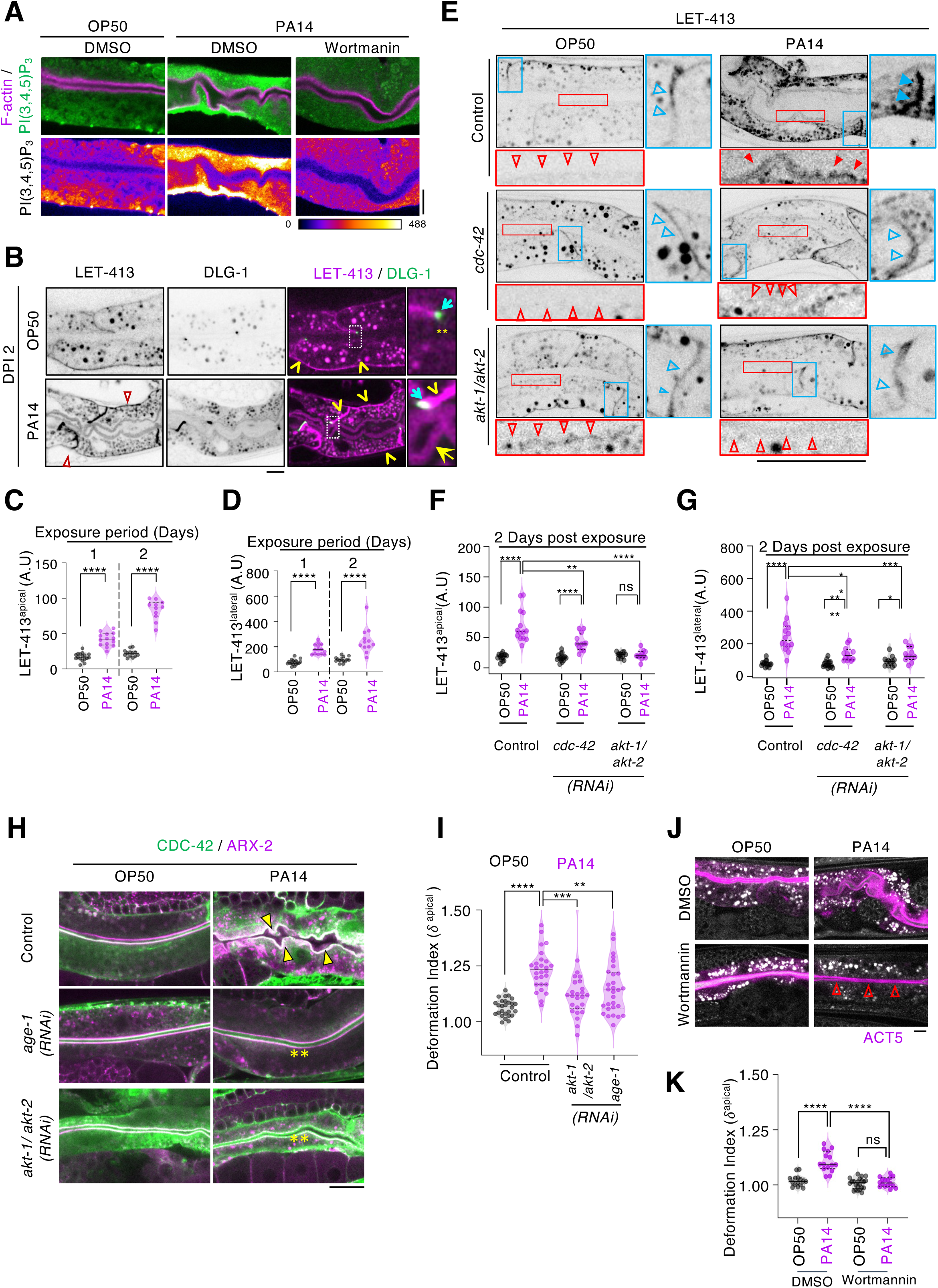
PI3K-AKT signaling mediates *P. aeruginosa* induced disruption of epithelial polarity and apical surface deformation. **(A)** Medial confocal sections of immune-stained *C. elegans* intestines showing PI(3,4,5)P_3_ localization following two days of exposure to *E. coli* or *P. aeruginosa* in NGM plates supplemented with DMSO or wortmannin. The enhanced apical localization of PI(3,4,5)P_3_ following *P. aeruginosa* infection was attenuated by wortmannin treatment, as highlighted in the enlarged ROIs. **(B)** Medial confocal sections of intestines expressing LET-413::mCherry following exposure to *E. coli* or *P. aeruginosa*. Following *P. aeruginosa* exposure, LET-413::mCherry signal increased along the apical membrane (yellow arrow) and lateral membrane (yellow arrowhead). DLG-1 marks cell junctions (cyan arrows). Yellow asterisks indicate the absence of detectable LET-413 signal from the apical surface in *C. elegans* exposed to *E. coli*. **(C, D)** Quantification of LET-413::mCherry fluorescence intensity at the (C) apical, and (D) lateral membranes following exposure to *E. coli* or *P. aeruginosa*. Plots show the median and interquartile range of each dataset. Each data point represents an individual worm. (n≥12). **(E)** Medial Confocal sections showing LET-413::mCherry localization in control (L4440), *cdc-42(RNAi),* and *akt-1/akt-2(RNAi)* animals following two days of exposure to *E. coli* or *P. aeruginosa*. Control animals exhibit infection-induced accumulation of LET-413::mCherry at the apical and basal regions, which is markedly reduced in *cdc-42(RNAi)* and *akt-1/akt-2(RNAi)* animals. Red outlined, unfilled arrowheads indicate reduced apical LET-413::mCherry accumulation. (F and G) Quantification of LET-413::mCherry fluorescence intensity at the (F) apical, and (G) lateral membranes in control, *cdc-42* and *akt-1/akt-2* (*RNAi*) animals following exposure to *E. coli* or *P. aeruginosa*. Violin plots show the median and interquartile range of each dataset. Each data point represents an individual worm. (n≥12). **(H)** Medial Confocal sections of C. elegans intestines co-expressing CDC-42 and ARX-2::TagRFP following RNAi mediated depletion of AGE-1 or AKT-1/AKT-2 and subsequent exposure to *E. coli* or *P. aeruginosa* for two days. Attenuation of PI3K-AKT signalling reduced *P. aeruginosa*-induced redistribution of ARX-2 and suppressed intestinal deformation. Yellow filed arrowheads indicate regions of intestinal deformation, whereas yellow asterisks denote suppression of deformation. **(I)** Violin plots showing quantification of the apical deformation index (*δ* ^apical^) in *age-1(RNAi) akt-1/akt-2(RNAi)* animals following two days of exposure to *E. coli* or *P. aeruginosa*. Plot shows the median and interquartile range of each data set (N =2, n≥26). **(J)** Representative medial confocal sections of *C. elegans* intestines expressing mCherry::ACT-5 following exposure to *E. coli* or *P. aeruginosa* in the presence of DMSO or wortmannin. Red outlined unfilled arrowheads indicate suppression of intestinal deformation following wortmannin treatment in *P. aeruginosa*-exposed worms. **(K)** Quantification of the apical deformation index (*δ* ^apical^) in worms treated with wortmannin and subsequently exposed to *E. coli* or *P. aeruginosa* exposure (n≥15). Scale bar in 20 µm. Statistical significance was determined using Mann -Whitney U test. ****p<0.0001, ***p <0.001, and **p <0.01.

We further validated these observations, by examining the localization of LET-413, the *C. elegans* Scribble homolog, and the CeAJ marker, DLG-1. The LET-413 complex maintain basolateral membrane identity by regulating the localization of apical polarity components^17,59^. In *E. coli*-fed worms, LET-413 localization was restricted to the basolateral membrane while the apical surface was largely devoid of LET-413 (Fig. 6B; OP50 panel, basolateral surface-yellow arrows, apical surface-yellow asterisks)^18^. In contrast, exposure to *P. aeruginosa* resulted in a significantly increased LET-413 localization along both the apical and lateral membranes (Fig. 6B; PA14 panel, apical surface - yellow arrows, lateral membrane-yellow open arrowhead, 6C and 6D), indicating disruption of epithelial polarity. Although cell junctions, marked by DLG-1, appeared extended, upon examining the single plane and projections of z-stack images, we noted that the apparent enlargement was due to the distortion of CeAJ resulting from apical surface deformation (PA14 Fig. 6B; PA14 panel - cyan arrows and S6C; yellow arrows). Whereas the CeAJ position was not altered following *P. aeruginosa* infection, junctional DLG-1 fluorescence intensity was increased relative to *E. coli*-fed controls (Fig. S6D).

The absence of intestinal deformation in *cdc-42(RNAi)* animals prompted us to investigate whether CDC-42 also regulates apico-lateral polarity following *P. aeruginosa* exposure. To address this question, we examined the LET-413 localization in control (L4440) and *cdc-42(RNAi)* worms. Enhanced apical accumulation of LET-413 observed in *P. aeruginosa*-exposed control animals was significantly reduced in *cdc-42(RNAi)* worms (Fig. 6E, 6F and 6G). Conversely, LET-413-depletion neither rescued apical surface deformation, not had any significant effect the apical localization or altered cytoplasmic levels of CDC-42 or ARP-2/3, consistent with previous reports of LET-413 having minimal role in *P. aeruginosa*-induced disruption of epithelial polarity (Fig S6E, S6F, and S6G)^60^. Together, these findings suggest that *P. aeruginosa* exposure results in the apical surface acquiring basolateral membrane characteristics.

### PI3K-AKT signalling couples polarity disruption to ARP-2/3 redistribution and intestinal deformation following *P. aeruginosa* infection

Activation of PI3K-AKT signalling and CDC-42-dependent actin assembly at the apical surface have been shown to be critical for *P. aeruginosa* infection in MDCK cells^76,78,79^. Similarly, long-lived *C. elegans* mutants in PI3K-AKT pathway exhibit enhanced resistance to *P. aeruginosa*^78,80^. To investigate whether the cellular responses induced by *P. aeruginosa* are conserved between mammalian epithelium and *C. elegans* enterocytes, we carried out simultaneous depletion of AKT-1 and AKT-2 and examined the intestinal localization of LET-413 following bacterial exposure (Fig. 6E). Similar to *cdc-42(RNAi)* animals, depletion of AKT-1/AKT-2 prevented the infection-induced apico-basal accumulation of LET-413, indicating that PI3K-AKT signalling is required for the polarity defects triggered by *P. aeruginosa* (Fig. 6E, 6F and 6G). Additionally, downregulated PI3K-AKT signalling in animals depleted of AGE-1, AKT-1/AKT-2 or Wortmannin treatment resulted in reduced apical deformation following *P. aeruginosa* exposure (Fig. 6H, S6H, S6I, 6I, 6J and 6K). Consistent with this phenotype, the infection-induced increase in cytosolic ARX-2 levels was also significantly attenuated in these animals (Fig. S6J and S6L), however apical accumulation was suppressed only in *age-1(RNAi)* worms (Fig. S6K and S6M). In contrast, CDC-42 levels remained elevated following infection despite disruption of PI3K-AKT signalling (Fig. S6N – S6Q) suggesting that following *P. aeruginosa* exposure, PI3K-AKT signalling might regulate ARP-2/3 localization downstream or independently of CDC-42 accumulation.

Our results, combined with previous observations in MDCK cells^75^, support a model in which *P. aeruginosa*-induced PI3K activation promotes PI(3,4,5)P_3_ enrichment, disrupting lipid homeostasis, apico-basal polarity and apical ARP-2/3-depnedent actin assembly. These findings further highlight epithelial polarity disruption as an early and critical event in pathogenesis of *P. aeruginosa* infection and intestinal deformation.

### Extracellular localization of *Pseudomonas* is sufficient to induce intestinal deformation

*Pseudomonas aeruginosa* can function as both an extracellular and intracellular pathogen infecting a wide range of hosts^81^. In MDCK cells, transient activation of PI3K pathway promotes the acquisition of basolateral characteristics by the apical membrane, thereby facilitating *P. aeruginosa* internalization^76^. Electron microscopy studies of *P. aeruginosa*-infected *C. elegans* have likewise suggested that the bacterium can invade intestinal epithelial cells^51,82^. We therefore sought to determine whether the intestinal deformation observed during infection results from *P. aeruginosa* breaching the epithelial barrier.

To address this question, we performed confocal imaging of live worms infected with GFP-expressing *E. coli* (OP50-GFP) or *P. aeruginosa* (PA14-GFP). Unlike *E. coli*, which were in very few numbers even after two days of exposure, *P. aeruginosa* cells formed visible aggregates throughout the intestinal lumen as early as one-day post infection (Fig. 7A). Notably, apical surfaces adjacent to these bacterial aggregates exhibited mild contortions that progressed to pronounced deformation by day-2 (Fig. 7A; red filled triangles in the PA14 panels).

**Figure 7:**
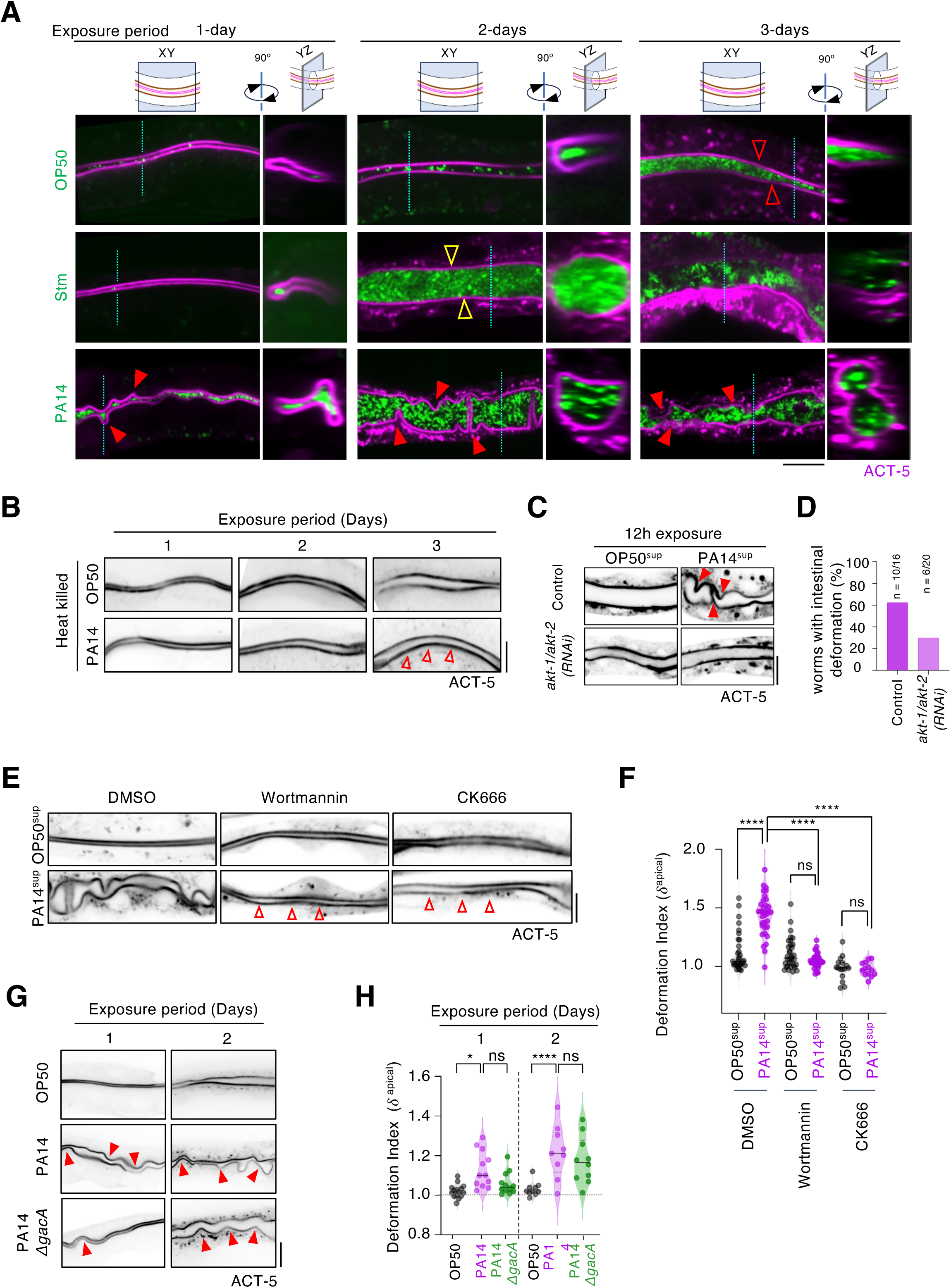
Secreted factors from extracellular *P. aeruginosa* induce intestinal deformation in *C. elegans*. **(A)** Confocal medial images of intestinal epithelium from *C. elegans* expressing ACT-::mCherry following three days of exposure to GFP-expressing *E. coli* (top panel), *S. typhymurium* (middle panel) and *P. aeruginosa* (bottom panel). Dashed lines indicate the plane used for orthogonal reconstruction, shown alongside the corresponding images. Red outlined, unfilled arrowheads (*E. coli*) and yellow outlined unfilled arrowheads (*S. typhimurium)* indicate intestinal distension (bloating), without apical surface deformation. Red filled arrowheads indicate apical surface deformation in *P. aeruginosa*-exposed animals. Orthogonal views show that bacteria remain confined to the intestinal lumen (N=2, n≥10). **(B)** Medial confocal images of intestines from *C. elegans* expressing mCherry::ACT-5 and maintained on heat-killed *E. coli* (top) or heat-killed *P. aeruginosa* (bottom). Red outlined, unfiled arrowheads indicate the absence of intestinal deformation following exposure to heat-killed *P. aeruginosa*. **(C)** Medial confocal sections of intestine in control (L4440) and *akt-1/akt-2(RNAi)* animals expressing mCherry::ACT-5 following 12 h exposure to cell-free, filter sterilized culture supernatants from *E. coli* or P*. aeruginosa*, as indicated. Red filed arrowheads indicate intestinal deformation induced by of *P. aeruginosa* culture supernatant. **(D)** Quantification of percentage of control and *akt-1/akt-2 (RNAi)* animals exhibiting intestinal deformation following *P. aeruginosa* exposure (n=16, control; n=20 *akt-1/akt-2 (RNAi)*. **(E)** Medial confocal sections of the intestine in worms expressing mCherry::ACT-5 treated with DMSO, wortmannin (1μM), or CK666(1μM), and exposed to cell free, filter sterilized culture supernatants from *E. coli* or *P. aeruginosa* for 12h. **(F)** Quantification of apical deformation index (*δ* ^apical^) in animals expressing intestinal mCherry::ACT-5 animals treated with DMSO, wortmannin (1µM) or CK666 (1µM) containing filter sterilized bacterial culture supernatants for 12 h (N=2, n≥15) **(G)** Medial confocal images of intestines from *C. elegans* expressing mCherry::ACT-5 following exposure to *E. coli* (OP50; top panel), wildtype *P. aeruginosa* (PA14; middle panel), or the *gacA* deletion mutant (PA14*ΔgacA*; bottom panel). Red filled arrowheads indicate regions of intestinal deformation. **(H)** Quantification of apical deformation index (*δ* ^apical^) in animals expressing intestinal mCherry::ACT-5 following exposure to *E. coli*, wild-type *P. aeruginosa,* or *P. aeruginosa ΔgacA*. Each data point represents individual animal and the violin plots indicate the median and interquartile of each data set. (n =10 per condition) Scale bar in 20 µm. Statistical significance was determined using Mann -Whitney U test. ****p<0.0001, ***p <0.001, and * p <0.05

**Figure 8:**
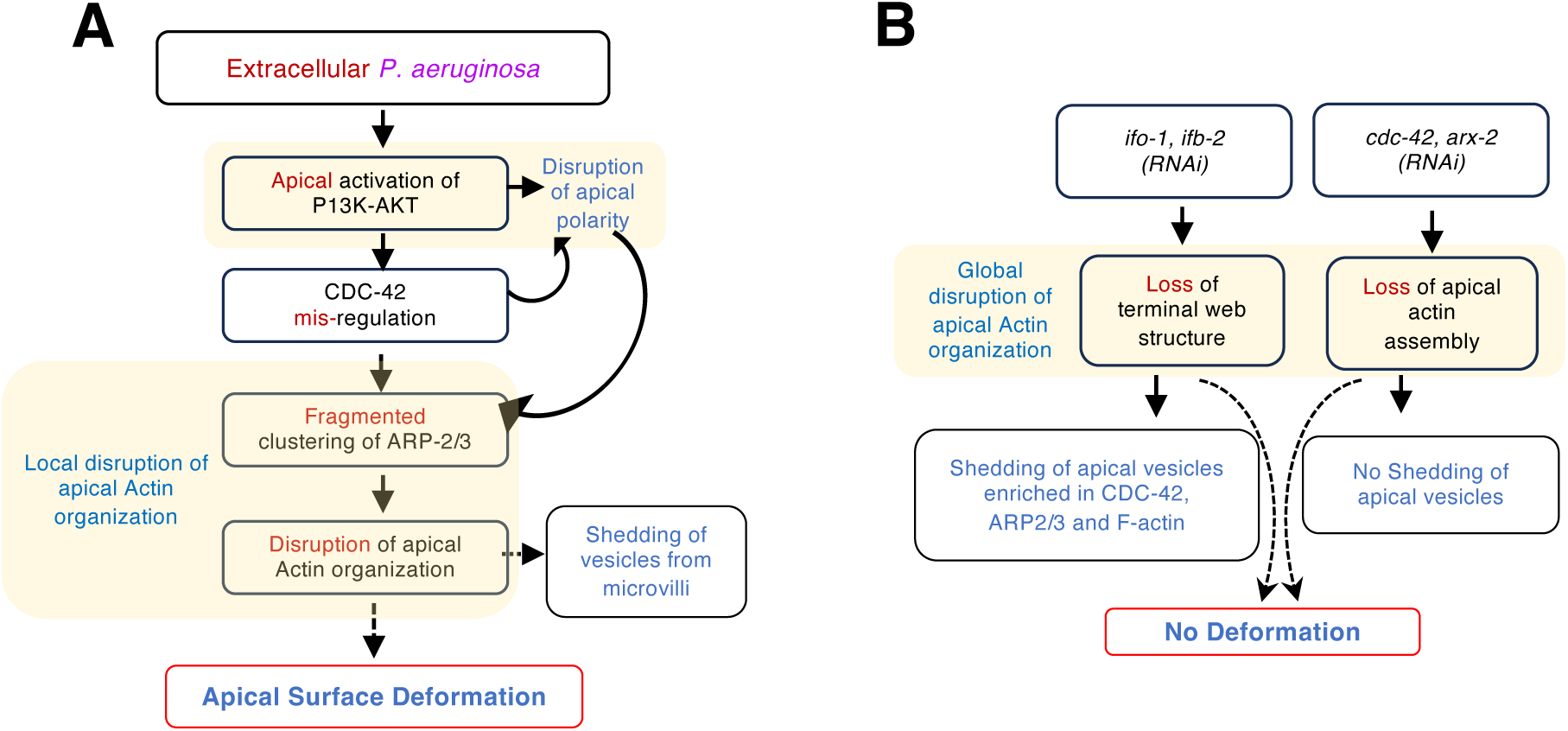
Mechanism of *P. aeruginosa*-induced intestinal deformation in *C. elegans*. **(A-B)** Schematic illustrating the proposed mechanism underlying *P. aeruginosa*-induced intestinal deformation in *C. elegans*.

However, orthogonal reconstruction (YZ plane) of confocal image stacks revealed that PA-GFP fluorescence remained confined to the intestinal lumen, with no detectable signal within the cytoplasm of intestinal cells (Fig.7A). Similar observations were obtained using lattice light-sheet microscopy, which likewise failed to detect intracellular bacteria (Fig. 4C). To further assess epithelial integrity, we examined the localization of CeAJ proteins, DLG-1 and α-catenin/HMP-1. In both *E. coli* and *P. aeruginosa*-infected animals, DLG-1 and HMP-1 localized as punctate apical structures in medial confocal sections and formed a continuous apical band in z-projections. Although infection-induced deformation resulted in a contorted pattern, the continuity of the junctional band was maintained (Fig. S6C and S7A). Consistent with this conclusion, propidium iodide was excluded from the intestinal cytoplasm and was confined to the lumen of infected animals, further confirming the absence of barrier disruption or widespread cellular invasion (Fig. S7B)^83^. These results suggest that epithelial barrier remained largely intact during deformation of the apical surface.

To determine whether apical deformation represents a general enterocyte response to intestinal pathogens, we performed parallel infection experiments using *Salmonella enterica serovar typhimurium* (Stm), a Gram-negative bacterium known to modulate CDC-42 activity during mammalian infections^84^ and cause lethality in *C. elegans (*Fig. S7C)^83,85^. Stm-GFP infection in worms expressing mCherry::ACT-5, resulted in robust intestinal colonization by day 2 similar to *P. aeruginosa* (Fig. S7D). A strong positive correlation between colony-forming units (CFU) counts and normalized lumenal bacterial fluorescence intensity further validated fluorescence-based measurements of intestinal colonization (Fig. S7D and S7E). Despite extensive colonization of the lumen, Stm-infected intestines did not exhibit any surface deformation (Fig. 7A; middle row, yellow open arrowheads). To exclude the possibility that reduced deformation in *cdc-42(RNAi)* animals resulted from differences in bacterial burden, we quantified the intestinal bacterial load and deformation for individual animals. Control and *cdc-42(RNAi)* worms exhibited similarly broad distributions of intestinal bacterial load; however, *cdc-42(RNAi*) animals consistently displayed markedly reduced intestinal deformation compared with controls (Fig. S7F). Thus, the lack of deformation following CDC-42 depletion cannot be explained by reduced bacterial colonization.

Together, these observations suggest that while intestinal distension is a general tissue response to microbial colonization of the lumen, mechanical deformation of apical surface is a specific response to *P. aeruginosa* exposure.

### Physical association between live *P .aeruginosa* and the intestinal surface is not essential for apical deformation

Physical interactions between *P. aeruginosa* and host cells are mediated by a variety of adhesins, including flagella, type IV pili, the type III secretion system (T3SS), and lectins^77,86,87^. In addition, several virulence factors, including bacterial pore-forming toxins (PFNs), have been implicated in remodelling the intestinal actin cytoskeleton ^88^. We therefore asked whether direct attachment of *P. aeruginosa* to the apical membrane of *C. elegans* intestine was necessary to trigger tissue deformation. First, we tested if exposure to live microbes is essential for inducing deformation of apical surface, by exposing animals to heat-killed *E. coli* or *P. aeruginosa* instead of live bacteria. Animals maintained on heat-killed bacteria as food source exhibited normal growth, maturation, and egg-laying, indicating that these food sources were not nutritionally limiting. However, unlike live *P. aeruginosa*, heat-killed *P. aeruginosa* failed to induce intestinal deformation (Fig. 7B; open red triangle). This observation is consistent with previous reports showing that heat-killed *P. aeruginosa* does not elicit immune responses in *C. elegans*^51^. Because extracellular *P. aeruginosa* induces intestinal deformation, we hypothesized that secreted factors to mediate this effect. To test this possibility, we exposed mCherry::ACT-5 expressing worms to cell-free, filter-sterilized supernatant collected from *P. aeruginosa* or *E. coli* cultures. Remarkably, 12-hour exposure to *P. aeruginosa* culture supernatant was sufficient to induce apical surface deformation in control intestines (Fig. 7C; red filled arrows in control panel, PA14^sup^). Inhibiting PI3K-AKT pathway or ARP2/3 mediated F-actin polymerization by RNAi or pharmacological drugs (using Wortmannin and CK666, respectively), significantly diminished intestinal deformation (Fig. 7C, 7D, 7E and 7F), suggesting that culture supernatant comprises of factor(s) that activate the same host signalling pathway as the intact bacteria.

The global response regulator, GacA governs the expression of numerous virulence factors secreted by *P. aeruginosa,* including toxins and biofilm regulators ^89–92^ and has also been implicated in inducing developmental defects and activating the unfolded protein response (UPR) in *C. elegans* intestinal epithelium ^42^. To assess the role of GacA in apical surface deformation, we exposed mCherry::ACT-5-expressing worms to *E. coli*, *P. aeruginosa* (PA14) or a *gacA* deletion mutant (PA14*1gacA*) strain. The intestines of worms exposed to PA14*1gacA* also exhibited intestinal deformation compared to those infected with wild-type PA14 (Fig. 7G and 7H). Importantly, a *gacA* deletion mutant (PA14*1gacA*) retained partial pathogenicity (Fig. S7G) ^93,94^. The persistence of deformation phenotype in the absence of GacA suggests the role of GacA-independent mechanism. Thus intestinal deformation is induced by heat-labile component(s), secreted into the lumen *P. aeruginosa* independent of GacA.

## DISCUSSION

Here we report that *P. aeruginosa* exposure induces two distinct morphological alterations in the *C. elegans* intestine; (a) lumenal distension, reflected by increased intestinal width, and (b) deformation of the apical surface, seen as wrinkling accompanied by release of membranous vesicles from the apical surface of the intestine. While lumen distension has been previously described as a general response to microbial colonisation, we show that apical surface deformation is specific to *P. aeruginosa* exposure.

In polarized MDCK cells, *P. aeruginosa* recruits PI3K to sites of bacterial attachment, leading to local enrichment of PI(3,4,5)P_3_, mis-localization of basolateral proteins to the apical membrane, and disruption of epithelial polarity through perturbation of Rho GTPases^66,67,75,95^. Likewise, *P. aeruginosa* infection in *C. elegans* led to apical PI(3,4,5)P_3_ enrichment and mis-localization of the basolateral marker LET-413, indicating a loss of apical membrane identity, though no apical markers were mis-localized basolaterally. Importantly, inhibition of either PI3K-AKT signalling or CDC-42-dependent actin assembly diminished both intestinal deformation and pathogen-induced lethality. Although LET-413 was mislocalized upon infection, depletion of LET-413 itself failed to rescue deformation, suggesting that the protective effect of CDC-42 depletion primarily reflects altered actin dynamics rather than restoration of epithelial polarity. RNAi mediated depletion of the JUN-kinase, KGB-1 did not suppress the intestinal deformation, discounting that actin phosphorylation as a possible cause for disruption in actin organization.

Consistent with this interpretation, depletion of ACT-5 or disruption of the ARP-2/3 complex phenocopied *cdc-42(RNAi)*, while attenuation of PI3K-AKT signalling limited ARP-2/3 distribution and prevented deformation. Together, these findings identify CDC-42-ARP-2/3-dependent actin remodelling as a key downstream effector of *P. aeruginosa-*induced enterocyte deformation. In uninfected animals, ARP-2/3 forms regularly spaced apical clusters associated with actin-binding proteins such as PLST-1 and FLN-2^72^. These clusters, as part of the terminal web support the organization of the apical F-actin network that stabilizes intestinal microvilli. Upon *P. aeruginosa* infection, ARP-2/3 clusters became fragmented leading to local disruption of apical actin organization. Intestinal apical actin organization has been shown to be extremely stable exhibiting limited turnover and mobility, therefore local disruptions in surface homogeneity may not be rapidly replenished, leading to progressive deterioration of the surface integrity ^21,72^(Fig. 4B). We propose that this fragmentation destabilizes the apical cortex and promotes shedding of actin-rich membrane material into the lumen. This model is consistent with electron microscopy studies reporting microvillar damage and host-derived debris in the *C. elegans* lumen during *P. aeruginosa* infection ^51,82^, as well as with observations of actin-rich protrusions in infected MDCK cells ^75^.

Interestingly, depletion of ARX-2 or ACT-5 prevented both vesicle release and surface deformation. Depletion of ACT-5 in *C. elegans* larval stages result in irregular and diminished microvillar structures, suggesting that ACT-5 is essential for the formation and maintenance of Intestinal microvilli^15,61^. While seeming counterintuitive, these results suggest that localized disruption of intact actin network and apical polarity promotes vesicle formation and surface deformation, whereas a uniform disruption as occurs following *act-5(RNAi)* or *arx-2(RNAi),* or pharmacological perturbation, does not. Similarly, Latrunculin A-mediated global actin depolymerization resulted in a 60-fold attenuation in *P. aeruginosa* infection in MDCK cells ^96^. We therefore speculate that spatial heterogeneity in cortical stability, rather than a general reduction in actin abundance, is an important determinant of apical surface deformation.

A notable feature of *P. aeruginosa* exposure was the accumulation of CDC-42, ARP-2/3, actin and PI(3,4,5)P_3_ within the intestinal cytoplasm. In polarized MDCK cells, *P. aeruginosa* attachment has been reported to increase apical CDC-42 levels and redistribute F-actin into the cytoplasm^96^. In contrast, although we observed cytoplasmic accumulation CDC-42, ARP-2/3 and actin in *C. elegans* enterocytes, their apical levels did not change. These observation suggest that the organization and regulation of the apical cytoskeleton in *C. elegans* intestine may differ from those in mammalian epithelial cell models, potentially contributing to distinct actin responses following *P. aeruginosa* infection. Our findings further suggest that the accumulated CDC-42 and ARP-2/3 pools do not promote F-actin polymerization, as the cytoplasmic actin is predominantly G-actin. Moreover, inhibition of PI3K-AKT signalling supressed ARP-2/3 redistribution without substantially affecting CDC-42 accumulation, indicating that increased CDC-42 abundance alone is insufficient to drive deformation. Whether cytoplasmic accumulation of PI(3,4,5)P_3_, polarity regulators and cytoskeletal components is sensed by the host as a pattern of pathogenesis remains to be addressed. In addition, MAPK signalling remained intact in *cdc-42(RNAi)* animals following infection, indicating that CDC-42 depletion does not impair microbial sensing. Together our findings support a model in which factors secreted by *P. aeruginosa* activate PI3K-AKT signalling at the apical membrane, promoting PI(3,4,5)P_3_ accumulation, disruption of apical membrane identity, fragmentation of ARP-2/3 organization, and destabilization of the microvillar actin network. This promotes shedding of microvillar material into the lumen and deformation of the intestinal surface.

The distinction between localized and a global disruption of the apical cytoskeleton is further illustrated by our genetic perturbations of actin and intermediate filament networks. Uniform disruption of F-actin organization through depletion of CDC-42, ARX-2, or ACT-5 did not itself induce deformation and instead protected against *P. aeruginosa*-induced damage. Similar to disruption of CDC-42-ARP-2/3-F-actin polymerization, attenuation of PI3K-AKT signalling or depletion of specific intermediate filament components suppressed pathogen-induced deformation. However, *ifo-1(RNAi)* and *ifb-2(RNAi)* animals released abundant luminal vesicles even in the absence of infection. Notably these vesicles contained apical determinants such as CDC-42, ARX-2 and F-actin, whereas vesicles released during *P. aeruginosa* infection were dominantly microvillar in composition and lacked detectable CDC-42 and ARX-2. These findings suggest that a pathogen-induced vesiculation occurs through a mechanism distinct from the generalized apical instability caused by disruption of intermediate filament network.

*P. aeruginosa* is generally considered as an extracellular pathogen, although intracellular infection has also been reported and may be associated with epithelial damage and loss of barrier integrity. Despite previous electron microscopy studies suggesting bacterial entry into *C. elegans* enterocytes ^51^, we found no detectable evidence of epithelial breach or intracellular colonization during the period examined. Instead, both confocal and lattice light sheet microscopy showed that bacteria remained confined to the intestinal lumen, while the integrity of the CeAJ was largely intact. Several factors may account for this difference. Our study focussed on the onset and progression of the intestinal deformation and intracellular infection may occur at a later stage than the period examined here. Differences in NGM composition (0.25% vs 0.35% peptone), culture conditions, or host physiology may also influence either bacterial virulence or host susceptibility and thereby affect the extent of intracellular invasion. Importantly our, data do not exclude the possibility that intracellular invasion occurs at later stages of infection or in a subset of cells below our detection threshold. Notably, exposure to cell-free *P. aeruginosa* culture supernatant but not heat-killed bacteria, reproduced intestinal surface deformation. These findings suggest that one or more secreted, heat-labile factor(s) mediate this response. One plausible candidate is outer membrane vesicles (OMVs), which carry diverse virulence-associated components, including toxins, quorum-sensing molecules, lipopolysaccharides and secondary metabolite^97–100^. In other host-pathogen systems, pore forming toxins (PFTs), produced by pathogens such as *S. aureus* and *B. thuringiensis*, can induce shedding of host cell membrane material as part of a membrane-repair response^101,102^. The membrane-disrupting activity of *P. aeruginosa* PA14 has been reported to depend on bacterial attachment and dedicated secretion systems^103,104^. However, despite pronounced surface deformation and extracellular vesicle release, we detected no evidence of widespread epithelial barrier disruption following *P. aeruginosa* exposure. Moreover the ability of cell-free culture to induce deformation suggests that direct bacterial attachment is not required for this response.

Previous studies have shown that dietary differences between HT115(DE3) and OP50 can influence host physiology and alter responses to *P. aeruginosa* infection ^105,106^. We observed a modest but reproducible difference in intestinal deformation between animals maintained on OP50 and those maintained on HT115(DE3) carrying the L4440 plasmid before exposure to either *P. aeruginosa* or OP50. Animals maintained on HT115(DE3) exhibited slightly less intestinal deformation following *P. aeruginosa* exposure. Nevertheless, pharmacological inactivation of PI3K-AKT signalling or ARP-2/3 mediated F-actin polymerization using Wortmannin or CK666 respectively, suppressed intestinal deformation in animals maintained exclusively in OP50. These results support a role for these [pathways in the deformation phenotype that is independent of the bacterial diet used for RNAi experiments.

Collectively, our findings provide insight into a potentially conserved mechanism through which *P. aeruginosa*-induced polarity defects alter epithelial cytoskeletal organization and tissue architecture during extracellular infection. We hypothesize that pathogen-mediated disruption of epithelial polarity may impair the directed secretion of antimicrobial peptides, lectins, and other host defence molecules. Such impairment could weaken intestinal defence, facilitating bacterial proliferation in the lumen and potentially promoting pathogen transmission within the host population.

## MATERIALS AND METHODS

### Growth and maintenance of strains

*C. elegans* and bacterial strains used in this study are listed in Table S1. Unless otherwise stated, all *C. elegans* strains were maintained at 20°C on Nematode Growth Medium (NGM) agar plates seeded with *E. coli* OP50^63^.All bacterial cultures were grown in Luria - Bertani (LB) broth at 37°C at 180 rpm.

### Bacterial infection assays

Infection assays were performed as previously described^94^ with minor modifications. Briefly, 40 µl of overnight bacterial culture (OD600 0.8-1) was spread on regular NGM plates containing 0.25% peptone and incubated at 37°C for approximately 4 h, followed by 24 h at room temperature prior to use. Age-synchronized L4 larvae (30 – 50 worms per condition) were transferred onto infection plates and maintained at 25° C throughout the experiment. For ‘slow-killing’ assays (Fig. S1L-M), modified NGM plates containing 0.35% peptone were prepared as described previously^94^.

For the heat-killed bacterial assay a single bacterial colony was inoculated into 100ml LB broth and cultured overnight at 37° C with shaking. Cultures were pelleted by centrifugation and incubated at 98° C for 45 min. Heat-killed bacteria were concentrated and seeded onto the standard NGM plates. Complete inactivation was confirmed by the absence of colony formation following plating on LB agar.

### Life span assays

For each condition, 100 age-synchronized L4 larvae were transferred to infection plates and monitored at 24 h intervals. Worms failing to respond to gentle touch were scored as dead. Missing animals were censored on the day of disappearance. Survival curves were generated using the Kaplan-Meier method, and median survival (TD_50_) was calculated for each condition. Statistical significance between survival curves was assessed using the log-rank test. All survival assays were performed in two independent biological replicates consisting of 100 worms each.

### RNA interference

RNA interference (RNAi) was performed by bacterial feeding using *E. coli* strain HT115 (DE3) carrying the L4440-based RNAi constructs targeting the following genes: *ced-10*, *cdc-42*, *rho-1*, *arx-2*, *act-*5, let*-413, age-1, akt-1, akt-2,* and *kgb-1*. Bacterial cultures were grown in LB broth supplemented with ampicillin (100 μg/mL) and tetracycline (12.5 μg/mL) at 37° C and seeded onto NGM agar containing 1 mM isopropyl β-D-thiogalactoside (IPTG) and 100 μg/mL ampicillin as previously described ^107^. Unless otherwise indicated, RNAi cultures were used undiluted. For *arx-2(RNAi)* and *act-5(RNAi)*, cultures were diluted 1:1 with HT115 (DE3) carrying the empty L4440 vector to minimize developmental defects. HT115 (DE3) carrying the empty L4440 served as the RNAi control. Age-synchronized L2 larvae were transferred onto RNAi for and maintained for 48h at 20° C before being used in infection assays. The *kgb-1* RNAi construct was generated by cloning a genomic fragment amplified using primers APO522 and APO523 (Table S1) into the L4440 vector.

### Fluorescence Microscopy and Image analysis

Live imaging was performed using an Olympus IX83 inverted microscope equipped with a CSU-W1 spinning-disk confocal head (Yokogawa Corporation, Tokyo, Japan), 10X UPlanS Apo (NA 0.40), 40X UPlanS Apo (NA 1.25), 60X UPlanS Apo (NA 1.42) or 100X UPlanS Apo (NA 1.40) objectives, and controlled using CellSens (version 2.3). Fluorophores were excited using 488 nm (GFP, eGFP and NeonGreen), 561 nm (mCherry, RFP, mScarlet and, dsRED) and 647 nm laser lines. Z-stacks were acquired with step sizes of 0.3 μm (100x objective) or 0.5 μm (60x and 40x objectives). Prior to imaging, worms were washed thoroughly with M9, immobilized in 0.025% levamisole, and mounted on 10% agarose pads. Images were acquired using a Prime BSI sCMOS camera (Photometrics, Tucson, AZ) at 1024×1024 pixels with 2x binning. Images shown in Fig. 3A (ACT-5) were acquired using an EMCCD camera (Photometrics, Tucson, AZ) with 512×512 pixels. Laser intensities and exposure periods were maintained constant across all experiments. Image processing analysis was carried out using ImageJ, ZEN 3.9 (blue edition, Zeiss), Imaris Viewer and MS-Excel.

To quantify the fluorescence intensity at the apical surface and in the cytoplasm, rectangular regions of interest (ROIs; 5 μm × 10-20 μm) were positioned at three randomly selected sites on each side of the intestinal lumen in both anterior and posterior intestinal regions (Fig. S2C). Mean fluorescence intensities were measured for each ROI and averaged to obtain a single value per animal.

### Quantification of intestinal distension and deformation (*δ* ^apical^, *δ* ^basal^)

Intestinal distension was quantified by measuring lumen diameter at multiple positions along the anterior and posterior intestine. Measurements from individual animals were averaged to obtain a mean lumen diameter. To quantify intestinal deformation, apical (*δ* ^apical^) and (*δ* ^basal^) deformation indices were calculated as ratio of contour length ‘l’ of the membrane surface to the corresponding linear body-axis distance ‘L’ (Fig. 1E). Higher values indicate greater membrane deformation. Between 15 and 20 worms were imaged for each treatment.

### Phalloidin Staining

For whole worm phalloidin staining, animals were washed thoroughly with 1X PBS buffer and fixed in 4% formaldehyde for 15 min at room temperature followed by acetone fixation at −20° C for 10 min. Following fixation, worms were permeabilized in 1X PBS containing 0.5% Triton X-100 and incubated with Alexa fluor 647-conjugated phalloidin (1:400) and DAPI (1:1000) for 2 h at room temperature in the dark. Samples were washed three times with 1X PBS and mounted in 10µl Fluormount-G™ for imaging.

### Quantification of Worm Body Length

To account for differences in body length during the quantification of intestinal deformation, worm length was measured from the pharyngeal region to the anal region. Measurements were obtained for control and RNAi-treated worms (*ifo-1, ifb-2, ifp-1, ifc-2, ifd-2, age-1*, and *act-5)* post two days of exposure to OP50 or PA14.

### Immunofluorescence detection of PI(3,4,5)P3 in Intestine

Intestines were dissected from animals immobilized in 0.2 mM levamisole and fixed in 4% paraformaldehyde in 1× PBS for 10-15 min at room temperature. Samples were blocked and gently permeabilized in 1xPBS containing 1% BSA and 0.2% saponin for 30 min at 4°C. Dissected intestines were incubated overnight at 4°C with a mouse monoclonal anti-PI(3,4,5)P3 antibody prepared in 1x PBS containing 0.05% saponin. Following three washes in 1x PBS supplemented with 0.05% saponin, samples were incubated with Alexa Fluor 488-conjugated donkey anti-mouse IgG for 2 h at room temperature. Samples were washed three times in 1x PBS containing 0.05% saponin and mounted in Fluoromount-G™ for imaging.

### Lattice Light sheet Imaging

Lattice light-sheet imaging was performed using a Lattice Lightsheet 7 microscope (Zeiss, Germany) equipped with a 13.3x/0.4 NA illumination objective positioned at 30° relative to the coverslip and a 44.83x/1.0 NA water-immersion detection objective positioned at 60°. Automatic water immersion and Alvarez optics were used during image acquisition. Dual-color imaging was performed using 488-nm and 560-nm excitation lasers and two ORCA-fusion sCMOS cameras (Hamamatsu). Images were acquired with a 10-ms exposure and a voxel size of 0.145 µm x 0.145 µm x 0.145 µm. Raw datasets were processed using constrained iterative deconvolution algorithm implemented in ZEN Blue 3.9 (Zeiss).

### Quantification of extracellular vesicle size and ERM-1 thickness

To estimate extracellular vesicle diameter, fluorescence intensity profiles were extracted from line scans passing through the centre of individual vesicles. To quantify ERM-1 thickness, line scans were drawn perpendicular to the apical surface membrane. Fluorescence intensities were normalized to the maximum intensity of each profile, maximum intensity positions were aligned and plotted as a function of distance. Profiles were fitted with a one-dimensional gaussian function: (Intensity(x)=A*exp[-(x-x_0_)^2^/(2*σ*^2^)]), where (A) represents the peak intensity, (x_0_) the position of the peak, and ′*σ*’ the standard deviation of the distribution. The full width at half maximum (FWHM) was calculated from the fitted standard deviation using the relationship FWHM = 2.355*σ*. Gaussian fitting and statistical analyses were performed in Python using the SciPy and NumPy libraries. Goodness-of-fit was assessed using the coefficient of determination (R²)., Mean FWHM, standard deviation (SD), and 95% confidence intervals (CI) were calculated from individual measurements. Distribution of vesicle widths / ERM-1 thickness was visualized using GraphPad Prism.

### Bacterial supernatant assay

Overnight cultures of *E. coli* and *P. aeruginosa* were centrifuged at 10K rpm for 10 min, and supernatants were passed through a 0.22 µm syringe filter to remove bacterial cells. Sterile supernatants were diluted 1:1 with S-basal buffer. Age-synchronized L1 larvae were subjected to either control (L4440) or *akt-1/akt-2 (RNAi)* treatment as described above. Following RNAi treatment, animals were washed extensively with M9 buffer and incubated in bacterial supernatants supplemented with S-basal buffer for 12 h at 25° C with gentle agitation. Worms were subsequently mounted on agarose pads and analyzed by fluorescence microscopy.

### Pharmacological Inhibition Assay

Pharmacological inhibition was performed using two approaches: (i) Inhibitor treatment on NGM plates and (ii) Inhibitor treatment in cell-free bacterial culture supernatants. Inhibitor treatment on NGM plates: CK-666 (1 µM), wortmannin (1 µM), latrunculin A (1.2 µM) or DMSO were spread NGM plates and incubated in dark at room temperature overnight to allow the inhibitors to diffuse and equilibrate through the agar. PA14 or OP50 cultures were subsequently spotted onto the plates and allowed to dry. Approximately 100 thoroughly washed, synchronized L4-stage animals were pre-incubated in M9 buffer containing the respective inhibitor or DMSO for 4 h (CK-666 and latrunculin A) or 6 h (wortmannin) at 25°C in the dark, with gentle agitation to maintain uniform suspension and adequate oxygenation. Following pre-incubation, the worms were transferred to the corresponding inhibitor-containing NGM plates seeded with PA14 or OP50 and maintained at 25 °C. The animals were subsequently harvested and processed for confocal imaging or immunofluorescence. (ii) Treatment with cell-free bacterial culture supernatants: Synchronized L4-stage worms were thoroughly washed, and approximately 100 animals were incubated in 200 µL of buffer containing CK-666 (1 µM), wortmannin (1 µM), or latrunculin A (1.2 µM), or an equivalent volume of DMSO as a control, for 6 h at 25 °C in the dark, with gentle agitation. Following pre-incubation, the animals were transferred to cell-free bacterial culture supernatants derived from PA14 or OP50 and supplemented with the respective inhibitor or DMSO. The worms were maintained under the same conditions for 12 h before being processed for confocal imaging.

### Intestinal barrier integrity assay

Intestinal integrity was assessed using propidium iodide (PI) (Sigma, P4170) uptake/exclusion assay as described previously ^108^. Worms exposed to *E. coli* or *P. aeruginosa* for two days were washed extensively with M9 buffer and incubated in M9 containing PI at 20 µg/ml for 45 minutes at 20°C with gentle agitation. Following staining, animals were washed twice with M9 buffer to remove excess dye and mounted on 10% agarose pads for imaging.

### Quantification of intestinal Bacterial Load

Intestinal bacterial load was estimated by determining colony-forming units (CFUs) per animal as described previously with minor modifications^11^. Briefly, ten worms were collected at 1, 2, or 3 days following exposure to GFP-expressing bacteria. Animals were washed extensively with 1x PBS containing 0.5 % levamisole for 10 min. Individual samples were homogenized in Subsequently the animals were homogenized in 50µl /1X PBS containing 0.01% Triton X-100. Homogenates were serially diluted and plated onto LB media containing the appropriate antibiotics. Following overnight incubation at 37°C, bacterial colonies were counted and CFUs per animal were calculated. Experiments were performed in three independent biological replicates.

### Quantification of bacterial fluorescence intensity

Normalized bacterial density within the intestinal lumen was estimated using ZEISS arivis Pro software (version 4.4). Approximately one-third of the image dataset was manually annotated and used to train a deep-learning segmentation model. Three classes were defined during training: intestinal lumen, bacteria, and background. The trained model was subsequently applied to the entire dataset using Deep Learning Segmenter module. Bacterial objects were segmented either using the trained deep-learning model or a blob-detection algorithm restricted to segmented intestinal lumen. Normalized bacterial fluorescence intensity for each worm was calculated by dividing the sum of intensity of segmented bacterial objects by the segmented intestinal lumen volume.

### Statistics

Statistical analysis were performed using the GraphPad Prism, version 6 unless otherwise stated. Data distributions were analysed using non-parametric statistical tests. Comparisons between two groups were performed using the Mann-Whitney U test. Survival curves were analyzed using the log-rank(Mantel-Cox) text. Correlation analyses were performed in Python using Spearman’s rank correlation coefficient.

## Data Availability Statement

Raw data underlying figures and analyses presented in this study are provided with the manuscript. Reagents generated in this study are available from the corresponding author upon request. Custom Scripts used for bacterial fluorescence quantification and correlation analyses are available at https://doi.org/10.5281/zenodo.20580797.

## ACKNOWLEDGEMENTS

This work was supported by DBT-Wellcome India Alliance Fellowship (IA/I/18/1/503624) and ANRF Core research grant (CRG/2023/004638) to A.P, Department of Biotechnology Junior research Fellowship (DBT/2024-25/AshokaUni/2486) to T.S., and core funding support from the Trivedi School of Biosciences, Ashoka University. We acknowledge the infrastructure support from the Central Bio-imaging facility and Ashoka-Zeiss Core Imaging Facility as well as Research Development Office (RDO) and Ashoka Global research Alliance (AGRA) at Ashoka University for facilitating various aspects of project implementation. We thank Ms. Megha Rai for help with vesicle size quantification, Ms. Shraddha Nirmal for technical assistance and Ms. Purna Pardeshi for some of the illustrations. We are grateful to the labs of Arnab Mukhopadhyay, Anindya Ghosh-Roy, Emily Troemel, Grégoire Michaux, Jogender Singh, Mike Boxem, Mahak Sharma and Varsha Singh, Rama Akondy for reagents, *C. elegans* and bacterial strains. Some strains were provided by the CGC, which is funded by NIH Office of Research Infrastructure Programs (P40OD010440). We are grateful to Ishani Sharma, Saravanan Palani and Jogender Singh for their feedback on the manuscript.

## List of Supporting Documents

The following list of supporting information is auxiliary to the main manuscript

1. Figures – ‘S1_Fig.tif’ – ‘S8_Fig.tif’

2. Supplementary Figure Legends – ‘S1_Text’

3. List of Main Figure Titles – ‘S2-Text’

4. List of Supplementary Figure Titles – ‘S3-Text’

5. List of Key Resources – ‘S1_Table-.xlsx’

6. Compressed Raw data file – ‘S1_Dataset-Metafile’

## References

1. Stuart, L. M., Paquette, N. & Boyer, L. Effector-triggered versus pattern-triggered immunity: how animals sense pathogens. Nat. Rev. Immunol. 13, 199–206 (2013).

2. Tse-Kang, S., Wani, K. A. & Pukkila-Worley, R. Patterns of pathogenesis in innate immunity: insights from C. elegans. Nat. Rev. Immunol. 10.1038/s41577-025-01167-0 (2025) doi:10.1038/s41577-025-01167-0.

3. Vance, R. E., Isberg, R. R. & Portnoy, D. A. Patterns of Pathogenesis: Discrimination of Pathogenic and Nonpathogenic Microbes by the Innate Immune System. Cell Host Microbe 6, 10–21 (2009).

4. Berg, M. et al. Assembly of the *Caenorhabditis elegans* gut microbiota from diverse soil microbial environments. ISME J. 10, 1998–2009 (2016).

5. Schulenburg, H. & Félix, M.-A. The Natural Biotic Environment of *Caenorhabditis elegans*. Genetics 206, 55–86 (2017).

6. Gravato-Nobre, M. J. et al. Multiple genes affect sensitivity of Caenorhabditis elegans to the bacterial pathogen Microbacterium nematophilum. Genetics 171, 1033–1045 (2005).

7. Lei, M., Tan, Y., Tu, H. & Tan, W. Neuronal basis and diverse mechanisms of pathogen avoidance in Caenorhabditis elegans. Front. Immunol. 15, 1353747 (2024).

8. Melo, J. A. & Ruvkun, G. Inactivation of Conserved C. elegans Genes Engages Pathogen- and Xenobiotic-Associated Defenses. Cell 149, 452–466 (2012).

9. Pereira, A. G., Gracida, X., Kagias, K. & Zhang, Y. *C. elegans* aversive olfactory learning generates diverse intergenerational effects. J. Neurogenet. 34, 378–388 (2020).

10. Prakash, D. et al. 1-Undecene from *Pseudomonas aeruginosa* is an olfactory signal for flight-or-fight response in *Caenorhabditis elegans*. EMBO J. 40, e106938 (2021).

11. Singh, J. & Aballay, A. Intestinal infection regulates behavior and learning via neuroendocrine signaling. eLife 8, e50033 (2019).

12. Zhang, Y., Lu, H. & Bargmann, C. I. Pathogenic bacteria induce aversive olfactory learning in Caenorhabditis elegans. Nature 438, 179–184 (2005).

13. Dimov, I. & Maduro, M. F. The C. elegans intestine: organogenesis, digestion, and physiology. Cell Tissue Res. 377, 383–396 (2019).

14. McGhee, J. The C. elegans intestine. WormBook 10.1895/wormbook.1.133.1 (2007) doi:10.1895/wormbook.1.133.1.

15. MacQueen, A. J. et al. ACT-5 Is an Essential *Caenorhabditis elegans* Actin Required for Intestinal Microvilli Formation. Mol. Biol. Cell 16, 3247–3259 (2005).

16. Armenti, S. T. & Nance, J. Adherens junctions in C. elegans embryonic morphogenesis. Subcell. Biochem. 60, 279–299 (2012).

17. Bossinger, O., Fukushige, T., Claeys, M., Borgonie, G. & McGhee, J. D. The apical disposition of the Caenorhabditis elegans intestinal terminal web is maintained by LET-413. Dev. Biol. 268, 448–456 (2004).

18. Legouis, R. et al. LET-413 is a basolateral protein required for the assembly of adherens junctions in Caenorhabditis elegans. Nat. Cell Biol. 2, 415–422 (2000).

19. Winter, J. F. et al. Caenorhabditis elegans screen reveals role of PAR-5 in RAB-11-recycling endosome positioning and apicobasal cell polarity. Nat. Cell Biol. 14, 666–676 (2012).

20. Jaffe, A. B. & Hall, A. Rho GTPases: Biochemistry and biology. Annu. Rev. Cell Dev. Biol. 21, 247–269 (2005).

21. Bernadskaya, Y. Y., Patel, F. B., Hsu, H. T. & Soto, M. C. Arp2/3 promotes junction formation and maintenance in the Caenorhabditis elegans intestine by regulating membrane association of apical proteins. Mol. Biol. Cell 22, 2886–2899 (2011).

22. Ma, L., Rohatgi, R. & Kirschner, M. W. The Arp2/3 complex mediates actin polymerization induced by the small GTP-binding protein Cdc42. Proc. Natl. Acad. Sci. U. S. A. 95, 15362–15367 (1998).

23. Martin-Belmonte, F. et al. PTEN-Mediated Apical Segregation of Phosphoinositides Controls Epithelial Morphogenesis through Cdc42. Cell 128, 383–397 (2007).

24. Ramalho, J. J., Nicolle, O., Michaux, G. & Boxem, M. In vivo control of the ezrin/radixin/moesin protein ERM-1 in C. elegans. bioRxiv 1–56 (2020).

25. Shafaq-Zadah, M., Brocard, L., Solari, F. & Michaux, G. AP-1 is required for the maintenance of apico-basal polarity in the *C. elegans*intestine. Development 139, 2061–2070 (2012).

26. Croce, A. et al. A novel actin barbed-end-capping activity in EPS-8 regulates apical morphogenesis in intestinal cells of Caenorhabditis elegans. Nat. Cell Biol. 6, 1173–1179 (2004).

27. Carberry, K. et al. The novel intestinal filament organizer IFO-1 contributes to epithelial integrity in concert with ERM-1 and DLG-1. Development 139, 1851–1862 (2012).

28. Geisler, F. et al. Intestinal intermediate filament polypeptides in C. elegans: Common and isotype-specific contributions to intestinal ultrastructure and function. Sci. Rep. 10, 3142 (2020).

29. Karabinos, A., Schmidt, H., Harborth, J., Schnabel, R. & Weber, K. Essential roles for four cytoplasmic intermediate filament proteins in Caenorhabditis elegans development. Proc. Natl. Acad. Sci. U. S. A. 98, 7863–7868 (2001).

30. Grimm-Günter, E.-M. S. et al. Plastin 1 Binds to Keratin and Is Required for Terminal Web Assembly in the Intestinal Epithelium. Mol. Biol. Cell 20, 2549–2562 (2009).

31. Hüsken, K. et al. Maintenance of the intestinal tube in Caenorhabditis elegans: The role of the intermediate filament protein IFC-2. Differentiation 76, 881–896 (2008).

32. Jahnel, O., Hoffmann, B., Merkel, R., Bossinger, O. & Leube, R. E. Mechanical Probing of the Intermediate Filament-Rich Caenorhabditis Elegans Intestine. Methods in Enzymology vol. 568 (Elsevier Inc., 2016).

33. Ruch, T. R. & Engel, J. N. Targeting the Mucosal Barrier: How Pathogens Modulate the Cellular Polarity Network. Cold Spring Harb. Perspect. Biol. 9, a027953 (2017).

34. Bastounis, E. E., Radhakrishnan, P., Prinz, C. K. & Theriot, J. A. Mechanical Forces Govern Interactions of Host Cells with Intracellular Bacterial Pathogens. Microbiol. Mol. Biol. Rev. 86, e00094–20 (2022).

35. Colonne, P. M., Winchell, C. G. & Voth, D. E. Hijacking Host Cell Highways: Manipulation of the Host Actin Cytoskeleton by Obligate Intracellular Bacterial Pathogens. Front. Cell. Infect. Microbiol. 6, (2016).

36. Fleiszig, S. M. et al. Epithelial cell polarity affects susceptibility to Pseudomonas aeruginosa invasion and cytotoxicity. Infect. Immun. 65, 2861–2867 (1997).

37. Tran, C. S. et al. Host Cell Polarity Proteins Participate in Innate Immunity to Pseudomonas aeruginosa Infection. Cell Host Microbe 15, 636–643 (2014).

38. Balla, K. M. & Troemel, E. R. Caenorhabditis elegans as a model for intracellular pathogen infection. Cell. Microbiol. 15, 1313–1322 (2013).

39. Mahajan-Miklos, S., Tan, M.-W., Rahme, L. G. & Ausubel, F. M. Molecular Mechanisms of Bacterial Virulence Elucidated Using a Pseudomonas aeruginosa–Caenorhabditis elegans Pathogenesis Model. Cell 96, 47–56 (1999).

40. Tan, M. W., Mahajan-Miklos, S. & Ausubel, F. M. Killing of Caenorhabditis elegans by Pseudomonas aeruginosa used to model mammalian bacterial pathogenesis. Proc. Natl. Acad. Sci. U. S. A. 96, 715–720 (1999).

41. Kuss-Duerkop, S. K. & Keestra-Gounder, A. M. NOD1 and NOD2 Activation by Diverse Stimuli: a Possible Role for Sensing Pathogen-Induced Endoplasmic Reticulum Stress. Infect. Immun. 88, e00898–19 (2020).

42. Pellegrino, M. W. et al. Mitochondrial UPR-regulated innate immunity provides resistance to pathogen infection. Nature 516, 414–417 (2014).

43. Dunbar, T. L., Yan, Z., Balla, K. M., Smelkinson, M. G. & Troemel, E. R. C. elegans detects pathogen-induced translational inhibition to activate immune signaling. Cell Host Microbe 11, 375–386 (2012).

44. McEwan, D. L., Kirienko, N. V. & Ausubel, F. M. Host translational inhibition by Pseudomonas aeruginosa exotoxin A triggers an immune response in Caenorhabditis elegans. Cell Host Microbe 11, 364–374 (2012).

45. Bollen, D. P., Reddy, K. C., Lascarez-Lagunas, L. I., Kim, D. H. & Colaiácovo, M. P. Germline mitotic quiescence and cell death are induced in *Caenorhabditis elegans* by exposure to pathogenic *Pseudomonas aeruginosa*. GENETICS 226, iyad197 (2024).

46. Peterson, N. D. et al. Non-canonical pattern recognition of a pathogen-derived metabolite by a nuclear hormone receptor identifies virulent bacteria in C. elegans. Immunity 56, 768–782.e9 (2023).

47. Ewbank, J. Signaling in the immune response. WormBook 10.1895/wormbook.1.83.1 (2006) doi:10.1895/wormbook.1.83.1.

48. Mukhopadhyay, A., Oh, S. W. & Tissenbaum, H. A. Worming pathways to and from DAF-16/FOXO. Exp. Gerontol. 41, 928–934 (2006).

49. Balla, K. M. & Troemel, E. R. Caenorhabditis elegans as a model for intracellular pathogen infection. Cell. Microbiol. 15, 1313–1322 (2013).

50. Cohen, L. B. & Troemel, E. R. Microbial pathogenesis and host defense in the nematode C. elegans. Curr. Opin. Microbiol. 23, 94–101 (2015).

51. Irazoqui, J. E. et al. Distinct Pathogenesis and Host Responses during Infection of C. elegans by P. aeruginosa and S. aureus. PLoS Pathog. 6, e1000982 (2010).

52. Peterson, N. D. et al. Pathogen infection and cholesterol deficiency activate the C. elegans p38 immune pathway through a TIR-1/SARM1 phase transition. eLife 11, e74206 (2022).

53. Pukkila-Worley, R. & Ausubel, F. M. Immune defense mechanisms in the Caenorhabditis elegans intestinal epithelium. Curr. Opin. Immunol. 24, 3–9 (2012).

54. Filipowicz, A., Lalsiamthara, J. & Aballay, A. TRPM channels mediate learned pathogen avoidance following intestinal distention. eLife 10, e65935 (2021).

55. Kumar, S. et al. Lifespan Extension in C. elegans Caused by Bacterial Colonization of the Intestine and Subsequent Activation of an Innate Immune Response. Dev. Cell 49, 100–117.e6 (2019).

56. Singh, J. & Aballay, A. Microbial Colonization Activates an Immune Fight-and-Flight Response via Neuroendocrine Signaling. Dev. Cell 49, 89–99.e4 (2019).

57. Szumowski, S. C. et al. Small GTPases promote actin coat formation on microsporidian pathogens traversing the apical membrane of *Caenorhabditis elegans*intestinal cells: Actin coats on microsporidia-containing vesicles. Cell. Microbiol. 18, 30–45 (2016).

58. Egge, N. et al. Age-Onset Phosphorylation of a Minor Actin Variant Promotes Intestinal Barrier Dysfunction. Dev. Cell 51, 587–601.e7 (2019).

59. McMahon, L., Legouis, R., Vonesch, J. L. & Labouesse, M. Assembly of C. elegans apical junctions involves positioning and compaction by LET-413 and protein aggregation by the MAGUK protein DLG-1. J. Cell Sci. 114, 2265–2277 (2001).

60. Riga, A. et al. Caenorhabditis elegans LET-413 Scribble is essential in the epidermis for growth, viability, and directional outgrowth of epithelial seam cells. PLOS Genet. 17, e1009856 (2021).

61. Göbel, V., Barrett, P. L., Hall, D. H. & Fleming, J. T. Lumen morphogenesis in C. elegans requires the membrane-cytoskeleton linker erm-1. Dev. Cell 6, 865–873 (2004).

62. Sepers, J. J., Ramalho, J. J., Kroll, J. R., Schmidt, R. & Boxem, M. ERM-1 Phosphorylation and NRFL-1 Redundantly Control Lumen Formation in the C. elegans Intestine. Front. Cell Dev. Biol. 10, (2022).

63. Stiernagle, T. Maintenance of C. elegans. WormBook 10.1895/wormbook.1.101.1 (2006) doi:10.1895/wormbook.1.101.1.

64. Ann Mack, N. & Georgiou, M. The interdependence of the Rho GTPases and apicobasal cell polarity. Small GTPases 5, 37–41 (2014).

65. Cowell, B. A., Evans, D. J. & Fleiszig, S. M. J. Actin cytoskeleton disruption by ExoY and its effects on *Pseudomonas aeruginosa* invasion. FEMS Microbiol. Lett. 250, 71–76 (2005).

66. Kazmierczak, B. I. & Engel, J. N. *Pseudomonas aeruginosa* ExoT Acts In Vivo as a GTPase-Activating Protein for RhoA, Rac1, and Cdc42. Infect. Immun. 70, 2198–2205 (2002).

67. Krall, R., Schmidt, G., Aktories, K. & Barbieri, J. T. Pseudomonas aeruginosa ExoT is a Rho GTPase-activating protein. Infect. Immun. 68, 6066–6068 (2000).

68. Pederson, K. J., Vallis, A. J., Aktories, K., Frank, D. W. & Barbieri, J. T. The amino-terminal domain of *Pseudomonas aeruginosa* ExoS disrupts actin filaments via small-molecular-weight GTP-binding proteins. Mol. Microbiol. 32, 393–401 (1999).

69. Sun, J. & Barbieri, J. T. ExoS Rho GTPase-activating Protein Activity Stimulates Reorganization of the Actin Cytoskeleton through Rho GTPase Guanine Nucleotide Disassociation Inhibitor. J. Biol. Chem. 279, 42936–42944 (2004).

70. Neukomm, L. J., Zeng, S., Frei, A. P., Huegli, P. A. & Hengartner, M. O. Small GTPase CDC-42 promotes apoptotic cell corpse clearance in response to PAT-2 and CED-1 in C. elegans. Cell Death Differ. 21, 845–853 (2014).

71. Wu, D. et al. CED-10-WASP-Arp2/3 signaling axis regulates apoptotic cell corpse engulfment in C. elegans. Dev. Biol. 428, 215–223 (2017).

72. Bidaud-Meynard, A. et al. High-resolution dynamic mapping of the *C. elegans* intestinal brush border. Development 148, (2021).

73. Stutz, K., Kaech, A., Aebi, M., Künzler, M. & Hengartner, M. O. Disruption of the C. elegans intestinal brush border by the fungal lectin CCL2 phenocopies dietary lectin toxicity in mammals. PLoS ONE 10, 1–23 (2015).

74. Crawley, S. W., Mooseker, M. S. & Tyska, M. J. Shaping the intestinal brush border. J. Cell Biol. 207, 441–451 (2014).

75. Kierbel, A., et al. *Pseudomonas aeruginosa* exploits a PIP3-dependent pathway to transform apical into basolateral membrane. J. Cell Biol. 177, 21–27 (2007).

76. Kierbel, A., Gassama-Diagne, A., Mostov, K. & Engel, J. N. The Phosphoinositol-3-Kinase–Protein Kinase B/Akt Pathway Is Critical for *Pseudomonas aeruginosa* Strain PAK Internalization. Mol. Biol. Cell 16, 2577–2585 (2005).

77. Thuenauer, R. et al. The Lectin LecB Induces Patches with Basolateral Characteristics at the Apical Membrane to Promote Pseudomonas aeruginosa Host Cell Invasion. mBio 13, e00819–22 (2022).

78. Garsin, D. A. et al. Long-Lived *C. elegans daf-2* Mutants Are Resistant to Bacterial Pathogens. Science 300, 1921–1921 (2003).

79. Yang, H. W. et al. Cooperative Activation of PI3K by Ras and Rho Family Small GTPases. Mol. Cell 47, 281–290 (2012).

80. Evans, E. A., Chen, W. C. & Tan, M. The DAF-2 insulin-like signaling pathway independently regulates aging and immunity in *C. elegans*. Aging Cell 7, 879–893 (2008).

81. Resko, Z. J., Suhi, R. F., Thota, A. V. & Kroken, A. R. Evidence for intracellular *Pseudomonas aeruginosa*. J. Bacteriol. 206, (2024).

82. Xue, F. et al. New tools to monitor Pseudomonas aeruginosa infection and biofilms in vivo in C. elegans. Front. Cell. Infect. Microbiol. 14, (2024).

83. Desai, S. K., Padmanabhan, A., Harshe, S., Zaidel-Bar, R. & Kenney, L. J. Salmonella biofilms program innate immunity for persistence in Caenorhabditis elegans. Proc. Natl. Acad. Sci. U. S. A. 116, (2019).

84. Bandyopadhyay, S. et al. Salmonella engages CDC42 effector protein 1 for intracellular invasion. J. Cell. Physiol. 239, (2024).

85. Aballay, A., Yorgey, P. & Ausubel, F. M. Supplementary material Salmonella typhimurium proliferates and establishes a persistent infection in the intestine of Caenorhabditis elegans Alejandro Aballay, Peter Yorgey and Frederick M. Ausubel. 2000 (2000).

86. Comolli, J. C., Waite, L. L., Mostov, K. E. & Engel, J. N. Pili Binding to Asialo-GM1 on Epithelial Cells Can Mediate Cytotoxicity or Bacterial Internalization by *Pseudomonas aeruginosa*. Infect. Immun. 67, 3207–3214 (1999).

87. Bucior, I., Pielage, J. F. & Engel, J. N. Pseudomonas aeruginosa Pili and Flagella Mediate Distinct Binding and Signaling Events at the Apical and Basolateral Surface of Airway Epithelium. PLoS Pathog. 8, e1002616 (2012).

88. Sitaram, A., Yin, Y., Zamaitis, T., Zhang, B. & Aroian, R. V. A Caenorhabditis elegans nck-1 and filamentous actin-regulating protein pathway mediates a key cellular defense against bacterial pore-forming proteins. PLoS Pathog. 18, e1010656 (2022).

89. Barta, T. M., Kinscherf, T. G. & Willis, D. K. Regulation of tabtoxin production by the lemA gene in Pseudomonas syringae. J. Bacteriol. 174, 3021–3029 (1992).

90. Kitten, T., Kinscherf, T. G., McEvoy, J. L. & Willis, D. K. A newly identified regulator is required for virulence and toxin production in Pseudomonas syringae. Mol. Microbiol. 28, 917–929 (1998).

91. Parkins, M. D., Ceri, H. & Storey, D. G. Pseudomonas aeruginosa GacA, a factor in multihost virulence, is also essential for biofilm formation. Mol. Microbiol. 40, 1215–1226 (2001).

92. Reimmann, C. et al. The global activator GacA of Pseudomonas aeruginosa PAO positively controls the production of the autoinducer N-butyryl-homoserine lactone and the formation of the virulence factors pyocyanin, cyanide, and lipase. Mol. Microbiol. 24, 309–319 (1997).

93. Feinbaum, R. L. et al. Genome-Wide Identification of Pseudomonas aeruginosa Virulence-Related Genes Using a Caenorhabditis elegans Infection Model. PLoS Pathog. 8, e1002813 (2012).

94. Tan, M.-W., Rahme, L. G., Sternberg, J. A., Tompkins, R. G. & Ausubel, F. M. *Pseudomonas aeruginosa* killing of *Caenorhabditis elegans* used to identify *P. aeruginosa* virulence factors. Proc. Natl. Acad. Sci. 96, 2408–2413 (1999).

95. Aktories, K. Bacterial protein toxins that modify host regulatory GTPases. Nat. Rev. Microbiol. 9, 487–498 (2011).

96. Kazmierczak, B. I., Mostov, K. & Engel, J. N. Epithelial Cell Polarity Alters Rho-GTPase Responses to *Pseudomonas aeruginosa*. Mol. Biol. Cell 15, 411–419 (2004).

97. Bauman, S. J. & Kuehn, M. J. Pseudomonas aeruginosa vesicles associate with and are internalized by human lung epithelial cells. BMC Microbiol. 9, 26 (2009).

98. Bauman, S. J. & Kuehn, M. J. Purification of outer membrane vesicles from Pseudomonas aeruginosa and their activation of an IL-8 response. Microbes Infect. 8, 2400–2408 (2006).

99. Kuehn, M. J. & Kesty, N. C. Bacterial outer membrane vesicles and the host–pathogen interaction. Genes Dev. 19, 2645–2655 (2005).

100. Mashburn, L. M. & Whiteley, M. Membrane vesicles traffic signals and facilitate group activities in a prokaryote. Nature 437, 422–425 (2005).

101. Husmann, M. et al. Elimination of a bacterial pore-forming toxin by sequential endocytosis and exocytosis. FEBS Lett. 583, 337–344 (2009).

102. Los, F. C. O. et al. RAB-5- and RAB-11-dependent vesicle-trafficking pathways are required for plasma membrane repair after attack by bacterial pore-forming toxin. Cell Host Microbe 9, 147–157 (2011).

103. Basso, P. et al. Pseudomonas aeruginosa Pore-Forming Exolysin and Type IV Pili Cooperate To Induce Host Cell Lysis. mBio 8, e02250–16 (2017).

104. Soong, G., Parker, D., Magargee, M. & Prince, A. S. The Type III Toxins of *Pseudomonas aeruginosa* Disrupt Epithelial Barrier Function. J. Bacteriol. 190, 2814–2821 (2008).

105. Reinke, S. N., Hu, X., Sykes, B. D. & Lemire, B. D. Caenorhabditis elegans diet significantly aYects metabolic profile, mitochondrial DNA levels, lifespan and brood size. Mol. Genet. Metab. 100, 274–282 (2010).

106. Revtovich, A. V., Lee, R. & Kirienko, N. V. Interplay between mitochondria and diet mediates pathogen and stress resistance in Caenorhabditis elegans. PLOS Genet. 15, e1008011 (2019).

107. Kamath, R. Genome-wide RNAi screening in Caenorhabditis elegans. Methods 30, 313–321 (2003).

108. Estes, K. A., Szumowski, S. C. & Troemel, E. R. Non-lytic, actin-based exit of intracellular parasites from c. elegans intestinal cells. PLoS Pathog. 7, (2011).

