## Supplementary material for "*P. aeruginosa* disrupts ARP-2/3 mediated apical F-actin organization to induce intestinal deformation in *C. elegans*": Supplemtentary Figure Legends

**Supplementary Figure S1: *P. aeruginosa* induces progressive deformation of the intestinal apical surface diverse growth conditions.**

**(A)** Quantification of intestinal lumen diameter ($\mu$m) in mCherry::ACT-5-expressing worms following exposure to *E. coli* or *P. aeruginosa.* Scatter plots show mean ±95% confidence interval (C.I). Each dot represents an individual worm. (N=3, n≥31 for Days 1 and 3, n≥21 for Day 2).

**(B)** Representative medial confocal sections of *C. elegans* intestine expressing mCherry::ACT-5 after two days of exposure to *E. coli* or *P. aeruginosa* at 20°C or 25°C.

(C) Quantification of the apical deformation index ($\delta^{apical}$) followingr two days of exposure to *E. coli* or *P. aeruginosa* at 20°C or 25°C (n ≥ 15).

**(D)** Quantification of the apical deformation index ($\delta^{apical}$) in worms exposed to *E. coli* (OP50) and two *P. aeruginosa* strains (PA14 and PAO1). Violin plots show the median and interquartile range of (n≥20, N=2).

**(E)** Quantification of '$\delta^{apical}$' in Cherry::ACT-5 expressing worms transiently exposed to *P. aeruginosa* for 12 h and 24 h before transfer to *E. coli* for an additional two days. Violin plot show median and interquartile range (n≥8).

**(F)** Representative medial confocal sections of *C. elegans* intestines expressing the basolateral marker LET-413::mCherry following exposure to *E. coli* or *P. aeruginosa*. Red lines indicate the continuous, unperturbed basal membrane under both conditions.

**(G)** Violin plot quantifying basal deformation index ($\delta_{basal}$) corresponding to (E) is plotted depicting the median and interquartile range of each data set. Each dot represents an individual animal (n≥12).

**(H)** Representaive medial plane confocal sections of the intestine in male worms expressing mCherry::ACT-5 after two days of exposure to *E. coli* or *P. aeruginosa*. Closed red arrowheads indicate intestinal deformation.

**(I)** Quantification of apical deformation index ($\delta^{apical}$) in male worms following three days of exposure to *E. coli* and *P. aeruginosa* . Violin plots shows the median and interquartile range. Each dot represents single worm (n≥9, N=2).

**(J)** Representative confocal images of wild-type N2 worms stained with Alexa Fluor 647-conjugated phalloidin after two days of exposure to *E. coli* or *P. aeruginosa.* Red arrows indicate intestinal deformation, and red asterisks denote the germline (n=20).

**(K)** Representative medial confocal sections of the intestinal epithelium in worms expressing LifeAct::mRuby following two days of exposure to *E. coli* or *P. aeruginosa*.

**(L)** Representative medial confocal sections of the *C. elegans* intestine expressing mCherry::ACT-5 following slow-killing assays with *E. coli* or *P. aeruginosa*. Red arrows indicate intestinal deformation.

**(M)** Quantification of '$\delta^{apical}$' corresponding to (K). Violin plots show median and interquartile range of each dataset. Each dot represents an individual animal ($n \geq 6$).

Scale bar - 20 $\mu$m. Statistical significance was determined using Mann-Whitney U test. ****$p<0.0001$, ***$p<0.001$, **$p<0.01$, and * $p<0.05$.

**Supplementary Figure S2: Depletion of CDC-42 suppresses apical surface deformation without impairing the *P.* aeruginosa-induced immune response**

**(A, B)** Quantification of ‘$\delta^{apical}$’ in mCherry::ACT-5 expressing animals subjected to control (L4440), *rho-1(RNAi)* (A) or *ced-10(RNAi)* (B) followed by two days of exposure to *E. coli* and *P. aeruginosa*. Violin plots show the median and interquartile range. Each dot represents individual worm (*rho-1(RNAi)* - N=2, n≥5 ; *ced-10(RNAi* N=2, n=7).

**(C)** Representative confocal image of worms expressing the immune-response reporter $P_{irg\text{-}1}$::GFP subjected to *cdc-42* and control (L4440) following *E. coli* or *P. aeruginosa* for two days. Scale bar, 50 $\mu$m.

**(D)** Quantification of $P_{irg\text{-}1}$::GFP fluorescence intensity in control (L4440) and *cdc-42(RNAi)* worms following two days of exposure to *E. coli* and *P. aeruginosa*.

**(E)** Schematic illustrating the method used to measure apical and cytoplasmic fluorescence intensities.

**(F)** Quantification of cytoplasmic F-actin fluorescence intensity following two days of exposure to *E. coli* or *P. aeruginosa*. Violin plots show the median and interquartile range. Each dot represents an individual worm (n=11).

Statistical significance was determined using Mann-Whitney U test. ****$p<0.0001$, **$p<0.01$, and *$p<0.05$

**Supplementary Figure S3: CDC-42 is required for the cytoplasmic accumulation of ARP-2/3 and actin following *P. aeruginosa* exposur, whereas CDC-42 accumulation is independent of ARP-2/3 and actin**

**(A)** Representative confocal images of *C. elegans* intestines showing RNAi-mediated depletion of CDC-42, ARX-2 or ACT-5.. Corresponding bar graphs quantify the reduction in respective protein levels following RNAi treatment (n≥5 animals per RNAi treatment).

**(B)** Quantification of apical mCherry::ACT-5 fluorescence intensity in worms subjected to *cdc42(RNAi)* or control (L4440), followed by two days of exposure to *E. coli* or *P. aeruginosa*. Violin plots show the median and interquartile range. Each dot represents an individual worm (N=2, n≥11).

**(C)** Representative medial confocal sections of the intestinal lumen in worms expressing CDC42::GFP and ARX2::TagRFP following *cdc-42 (RNAi)* or control (L4440) and subsequent exposure to *E. coli* or *P. aeruginosa*. Images were acquired and displayed using identical intensity settings to visualize RNAi-mediated depletion. Yellow arrows indicate the intestinal deformation.

**(D, E)** Quantification of (D) apical and (E) cytoplasmic ARX-2::TagRFP fluorescence intensities in *cdc-42 (RNAi)* and control (L4440) animals (from Figure S3B), subjected to *E. coli* and *P. aeruginosa* exposure. Violin plot show the median and interquartile range (n≥11).

**(F, G)** Quantification of (F) apical , and (G) cytoplasmic CDC-42::GFP fluorescence intestines in *arx-2(RNAi)* and control (L4440) (from Figure 3C), following two days of exposure to *E. coli* or *P. aeruginosa*. Violin plots show the median and interquartile range (n≥8, N=2).

**(H, I)** Quantification of (H) cytoplasmic and (I) apical CDC42::GFP fluorescence intensities in *act-5(RNAi)* and control (L4440) animals (from Figure 3D), following two days of exposure to *E. coli* or *P. aeruginosa*. Violin plots show the median and interquartile range (n≥8).

**(J, K)** Quantification of (J) cytoplasmic and (K) apical ARX-2::TagRFP fluorescence intensities in *act-5(RNAi)* and control (L4440) animals (from Figure 3D), following exposure to *E. coli* or *P. aeruginosa*. Violin plots show the median and interquartile range of each dataset (n≥8).

Scale bar, 20*μ*m. Statistical significance was determined using Mann-Whitney U test. ****$p<0.0001$, ***$p<0.001$, **$p<0.01$, and * $p<0.05$

**Supplementary Figure S4: Lumenal vesicles released following *P. aeruginosa* infection contain an F-actin core**

**(A)** Line-scan fluorescence intensity profiles of mCherry::ACT5 and PGP-1::GFP across extracellular vesicles in worms following two days of exposure to *P. aeruginosa* (n=20 vesicles).

(**B)** Quantification of the full width half maximum (FWHM) of mCherry::ACT-5-positive extracellular vesicles following two days of exposure to *P. aeruginosa*. Violin plots show the median and interquartile range of each dataset (n=18 vesicles).

**(C)** Representative medial confocal sections of the intestinal lumen in worms expressing IFB-2::mScarlet following *kgb-1(RNAi)* or control (L4440) and subsequent exposure to *E. coli* or *P. aeruginosa* for two days (n≥10).

Scale bar, 20μm.

**Supplementary Figure S5:Effects of ARP-2/3 and intermediate filame deletiom on IFB-2 and ARX-2 following *P. aeruginosa* exposure**

**(A, B)** Quantification of (A) apical and (B) cytoplasmic IFB-2::mScarlet fluorescence intensities in control (L4440) or *arx-2(RNAi)* animals following one day of exposure to *E. coli* or *P. aeruginosa* . Violin plots show the median and interquartile range of each data set (N=2, n≥16).

**(C, D)** Quantification of (C) apical and (D) cytoplasmic ARX-2::tagRFP fluorescence intensities in control (L4440) or *ifo-1(RNAi)* animals following one day of exposure to *E. coli* or *P. aeruginosa* . Violin plots show the median and interquartile range of each data set (n≥16).

**(E)** Representative confocal images of the *C. elegans* intestines IFB-2::mScarlet in control(L4440) and ifb-2(RNAi) animals. The accompanying bar graph quantifies the reduction in IFB-2::mScarlet fluorescence intensity following the *ifb-2* RNAi (n≥5).

**(F)** Quantification of body length in control (L4440) and animals subjected to RNAi0mediated depletion of the intermediate filament proteins (IFO-1, IFB-2, IFC-2, IFP-1, and IFD-2), or of AGE-1, or ACT-5 following two days of exposure to *E. coli* or *P. aeruginosa*. Violin plots show the median and interquartile range of each dataset (n ≥ 12). Statistical significance was determined using one-way ANOVA.

Scale bar, 20$\mu$m. Statistical significance in (A) – (E) was determined using Mann-Whitney U test. ****p<0.0001, ***p <0.001, **p<0.01, and * p <0.05

**Supplementary Figure S6: Attenuation of PI3K-AKT signalling suppresses *P. aeruginosa*-induced increase in cytoplasmic ARX-2 levels**

(A) Quantification of PI(3,4,5)$P_3$ immunofluorescence intensities in the apical, cytoplasmic, and basal regions of the intestine in worms treated with DMSO or wortmannin before exposure to *P. aeruginosa* (n≥11).

**(B)** Representative confocal images of the *C. elegans* intestine expressing LET-413::mScarlet in control (L4440) and *let-413(RNAi)* animals. The accompanying bar graph quantifies the reduction in LET-413::mScarlet fluorescence intensity following RNAi treatment (n ≥ 5 animals).

**(C)** Representative medial confocal section and corresponding maximum-intensity z-projection of the intestine in worms expressing DLG-1::eGFP following exposure (2 days) to *E. coli* and *P. aeruginosa*. CeAJs appear continuous un-perturbed under both conditions .

**(D)** Quantification of DLG-1::eGFP fluoresence intensity from medial confocal sections (in Fig. S6C), following two days of exposure to *E. coli* or *P. aeruginosa.* Violin plots show the median and interquartile range of each dataset (n≥12).

**(E)** Medial confocal images of intestines from *let-413(RNAi)* and control (*L4440*) animals co-expressing CDC42::GFP and ARX2::TagRFP following two days of exposure to *E. coli* or *P. aeruginosa*.

**(F-G)** Quantification of (F) cytoplasmic CDC42::GFP and (G) cytoplasmic ARX-2::TagRFP fluorescence intensities in *let-413(RNAi)* animals following two days of exposure to *E. coli* or *P.* aeruginosa. Violin plots show the median and interquartile range of each dataset (n≥10)

**(H)** Quantification of '$\delta^{apical}$' in wild-type N2 and *age-2(yw23) I; age-1(hx546)* double-mutant animals following two days of exposure to *P. aeruginosa*. Violin plots show the median and interquartile range of each dataset (n≥11).

**(I)** Representative medial confocal sections of intestinal lumen of animals expressing IFB-2::mScarlet following control (*L4440*) and *akt-1/akt-2(RNAi)* and subsequent exposure to *E. coli* or *P. aeruginosa.*

**(J-K)** Quantification of cytoplasmic (J) and, apical (K) ARX2::TagRFP fluorescence intensities in *age-1(RNAi)* animals following two days of exposure to *E. coli* or *P. aeruginosa.* Violin plots show the median and interquartile range of each dataset (n≥17, N=2).

**(L-M)** Quantification of (L) cytoplasmic and (M) apical ARX2::TagRFP fluorescence intensities in *akt-1/akt-2(RNAi)* animals following two days of exposure to *E. coli* or *P. aeruginosa*. Violin plots show the median and interquartile range of each dataset). ($n \geq 20$, $N=2$).

**(N-O)** Quantification of (N) cytoplasmic and (O) apical CDC-42::GFP fluorescence intensities in *age-1(RNAi)* and control animals following two days of exposure to *E. coli* or *P. aeruginosa* exposure. Violin plots show the median and interquartile range of each dataset ($n \geq 17$, $N=2$).

**(P-Q)** Quantification of (P) cytoplasmic and (Q) apical CDC-42::GFP fluorescence intensity in *akt-1/akt-2 (RNAi)* and control animals following two days of exposure to *E. coli* or *P. aeruginosa*. Violin plots show the median and interquartile range of each dataset ($n \geq 20$, $N=2$).

Scale bar, 20 $\mu$m. Statistical significance was determined using Mann-Whitney U test. ****$p<0.0001$, ***$p<0.001$, **$p<0.01$, and * $p<0.05$

**Supplementary Figure S7: CeAJ and Intestinal barrier is not compromised following *P. aeruginosa* exposure**

**(A)** CeAJs are not disrupted upon *P. aeruginosa* exposure. Medial confocal sections and corresponding maximum-intensity z-projections of the intestines of worms expressing HMP-1::GFP following two days of exposure to *E. coli* or *P. aeruginosa*. (n≥10).

**(B)** Medial confocal sections and corresponding maximum intensity z- projections of the intestines of wildtype N2 worms treated with propidium iodide (PI) following two days of exposure to *E. coli* or *P. aeruginosa.* No detectable PI uptake into the intestinal epithelium was observed, consistent with the preservation of epithelial integrity (n≥10).

**(C)** Representative Kaplan-Meier survival curves of L4-stage worms exposed to *Salmonella enterica serovar Typhimurium* (SL1344) at $25^{0}$ C (n = 100 worms/condition, N=2).

**(D)** Quantification of colony-forming units (CFU) per animal in mCherry::ACT-5-expressing worms following three days of exposure to fluorescently labelled *E. coli, S. typhimurium* or *P. aeruginosa*. Scatter plots show the mean±SEM (n=10 per condition;. N=3 for *E. coli and P. aeruginosa*, and N = 2 for *S. typhimurium*).

**(E)** Correlation between CFUs per animal and luminal *P. aeruginosa* fluorescence intensity normalized to intestinal lumen volume on Days 1, 2, and 3 following exposure. Spearman correlation ($\rho$) = 1.000

**(F)** Correlation between '$\delta^{apical}$' and intestinal bacterial density in control (L4440) and *cdc-42(RNAi)* animals following two days of exposure to *P. aeruginosa.* Spearman correlation ($\rho$) for control (L4440) = -0.51; and *cdc-42(RNAi)* = -0.20, suggesting non-correlated. (n≥11)

**(G)** Representative Kaplan-Meier survival curves of animals described in Fig. 7G following exposure to *E. coli* (OP50), wild-type *P. aeruginosa* (PA14) or the *P. aeruginosa gacA* deletion mutant (*PA14ΔGacA*) (n = 50 worms per condition, N=2).
