## Supplementary Figures for "*P. aeruginosa* disrupts ARP-2/3 mediated apical F-actin organization to induce intestinal deformation in *C. elegans*"

### Supplementary Figure S1

*Divyashree et. al (2026)*

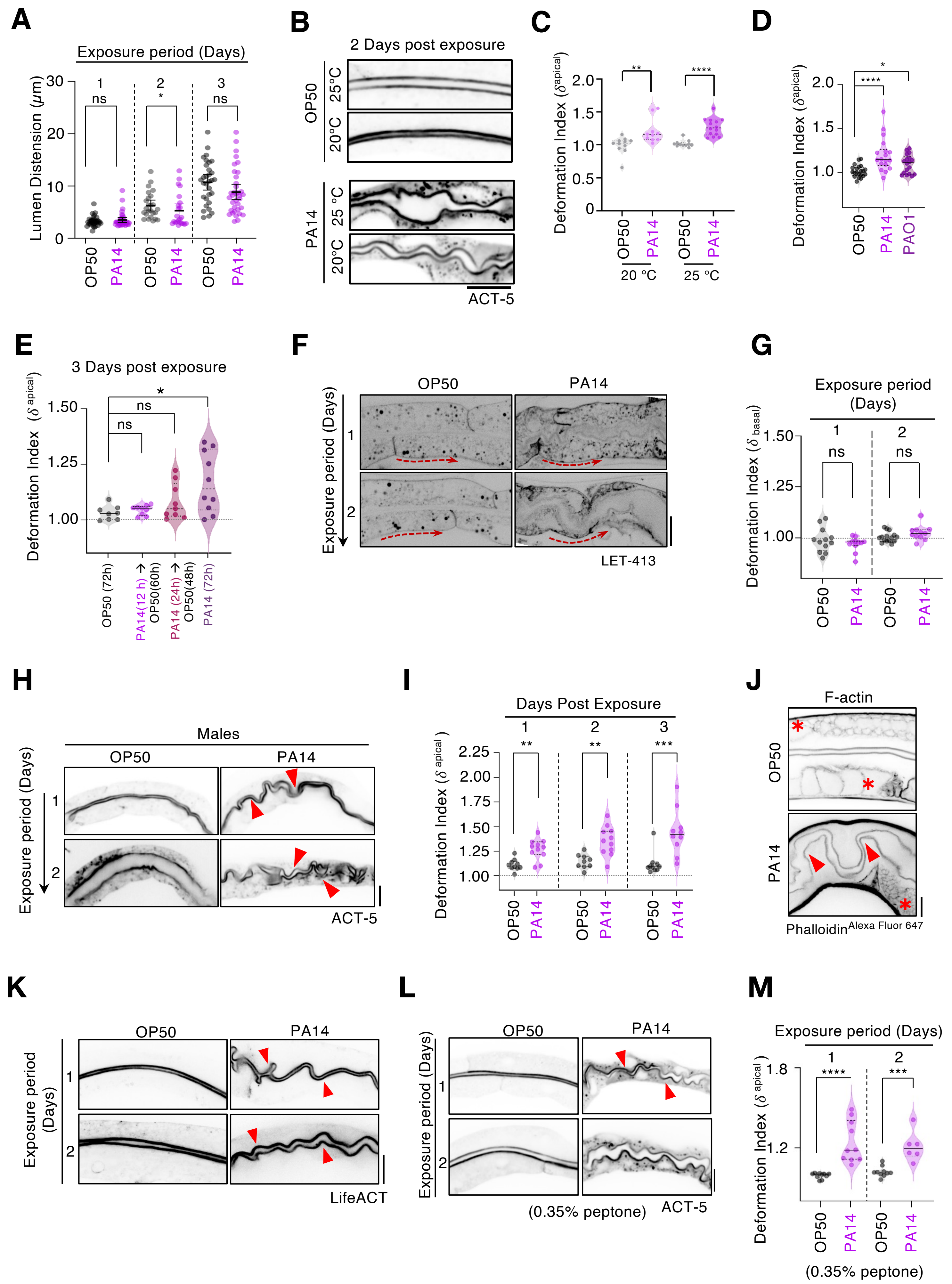

#### Supplementary Figure S2

*Divyashree et. al (2026)*

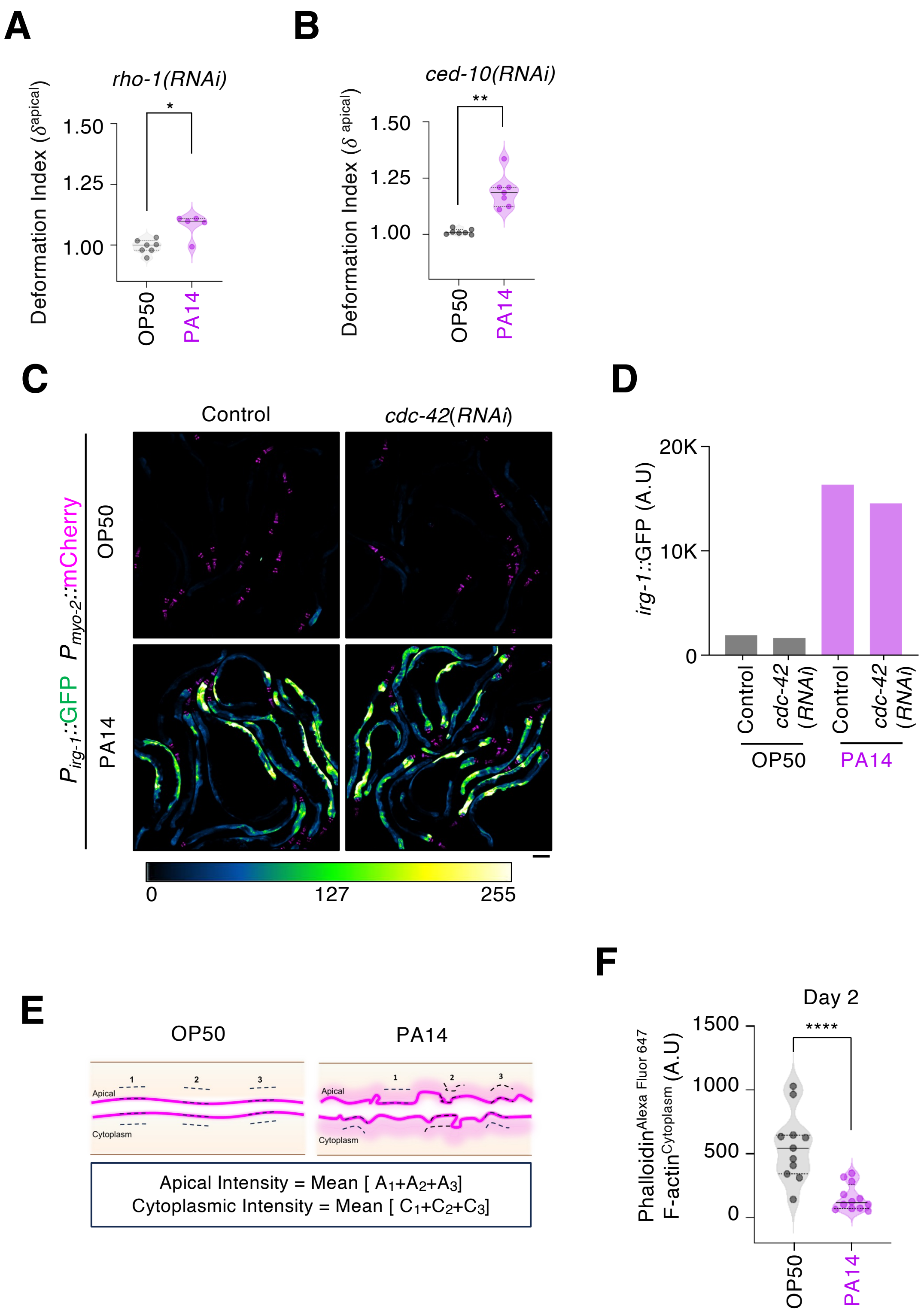

### Supplementary Figure S3

*Divyashree et. al (2026)*

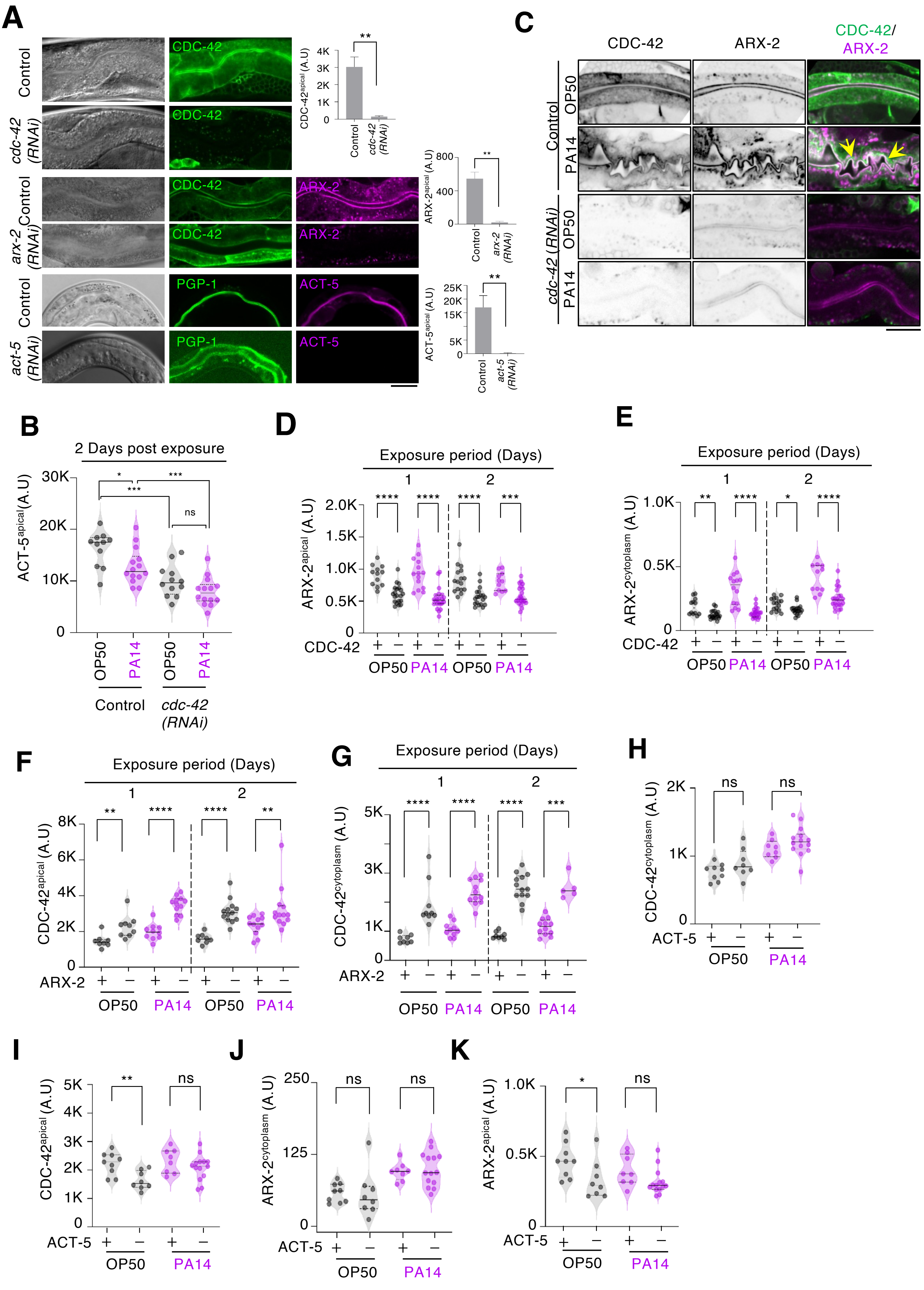

### Supplementary Figure S4

*Divyashree et. al (2026)*

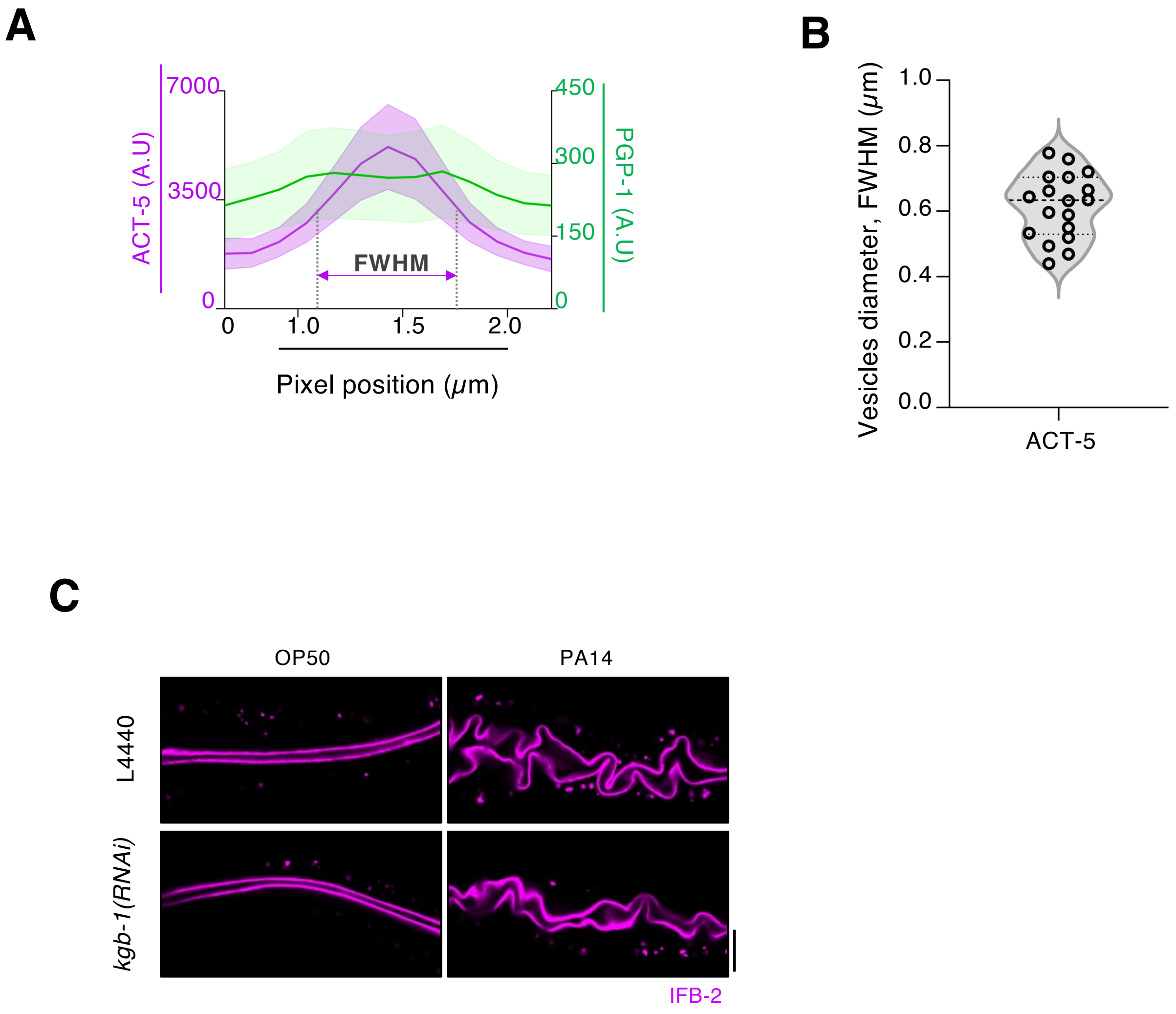

### Supplementary Figure S5

*Divyashree et. al (2026)*

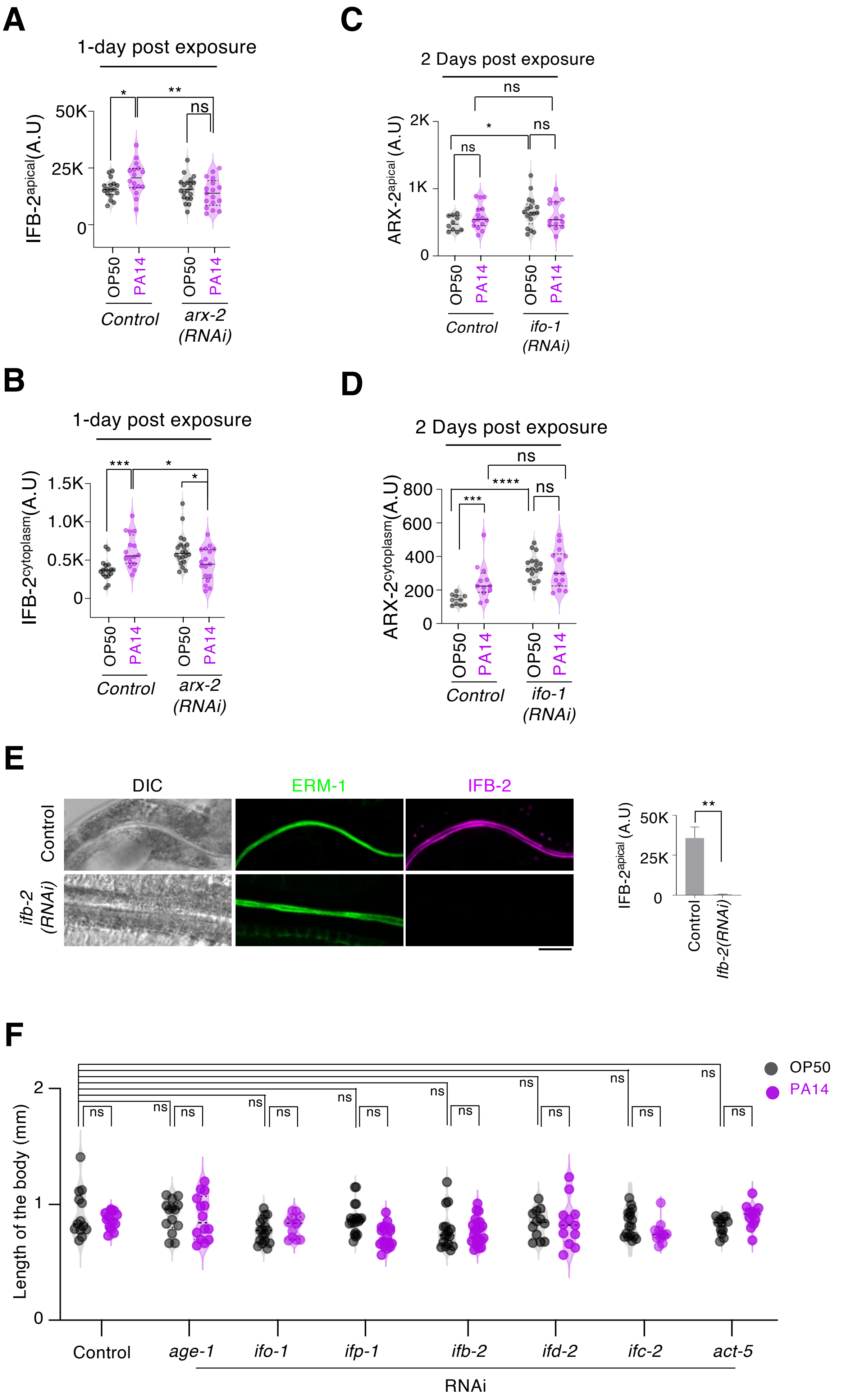

### Supplementary Figure S6

*Divyashree et. al (2026)*

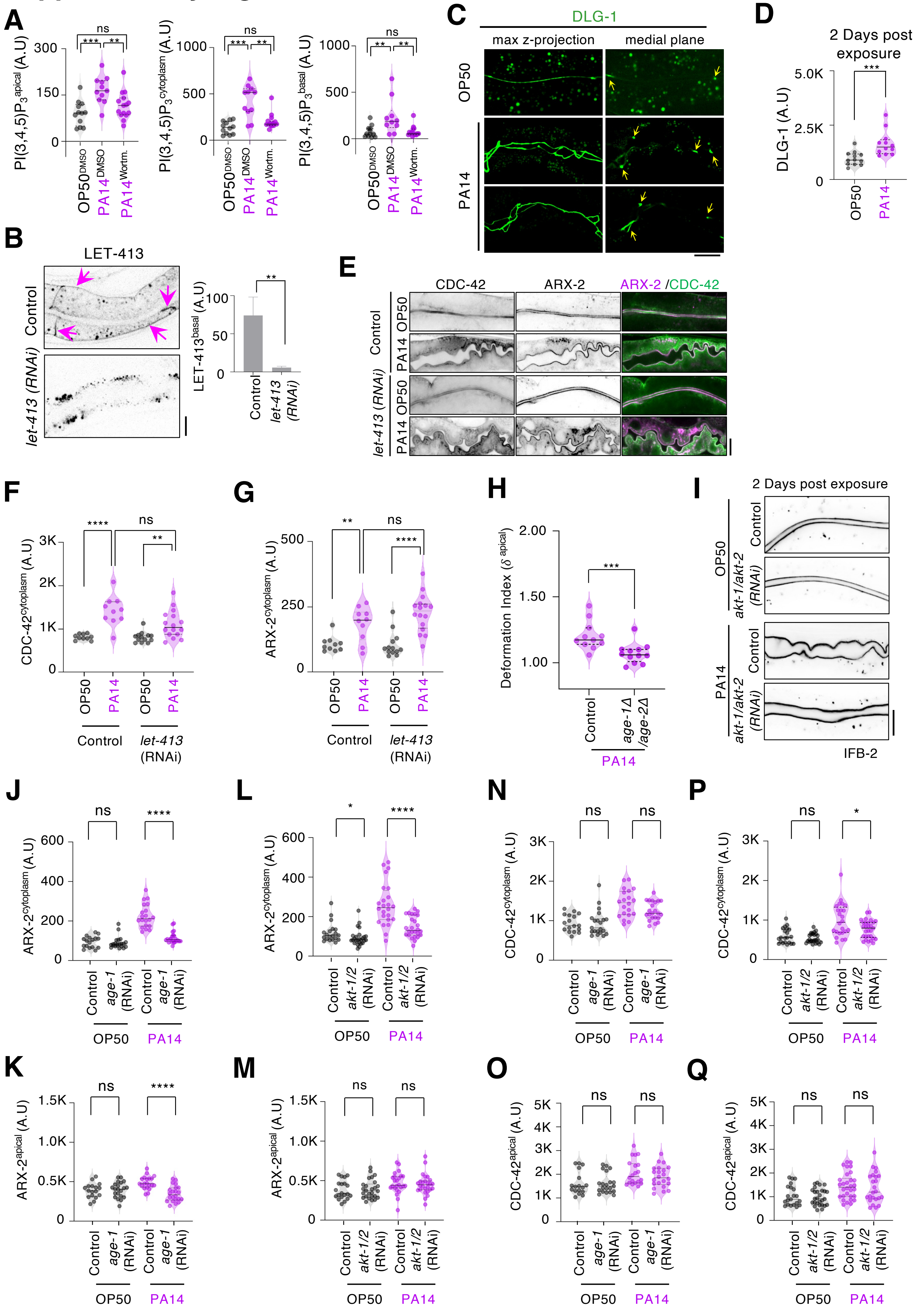

### Supplementary Figure S7

*Divyashree et. al (2026)*

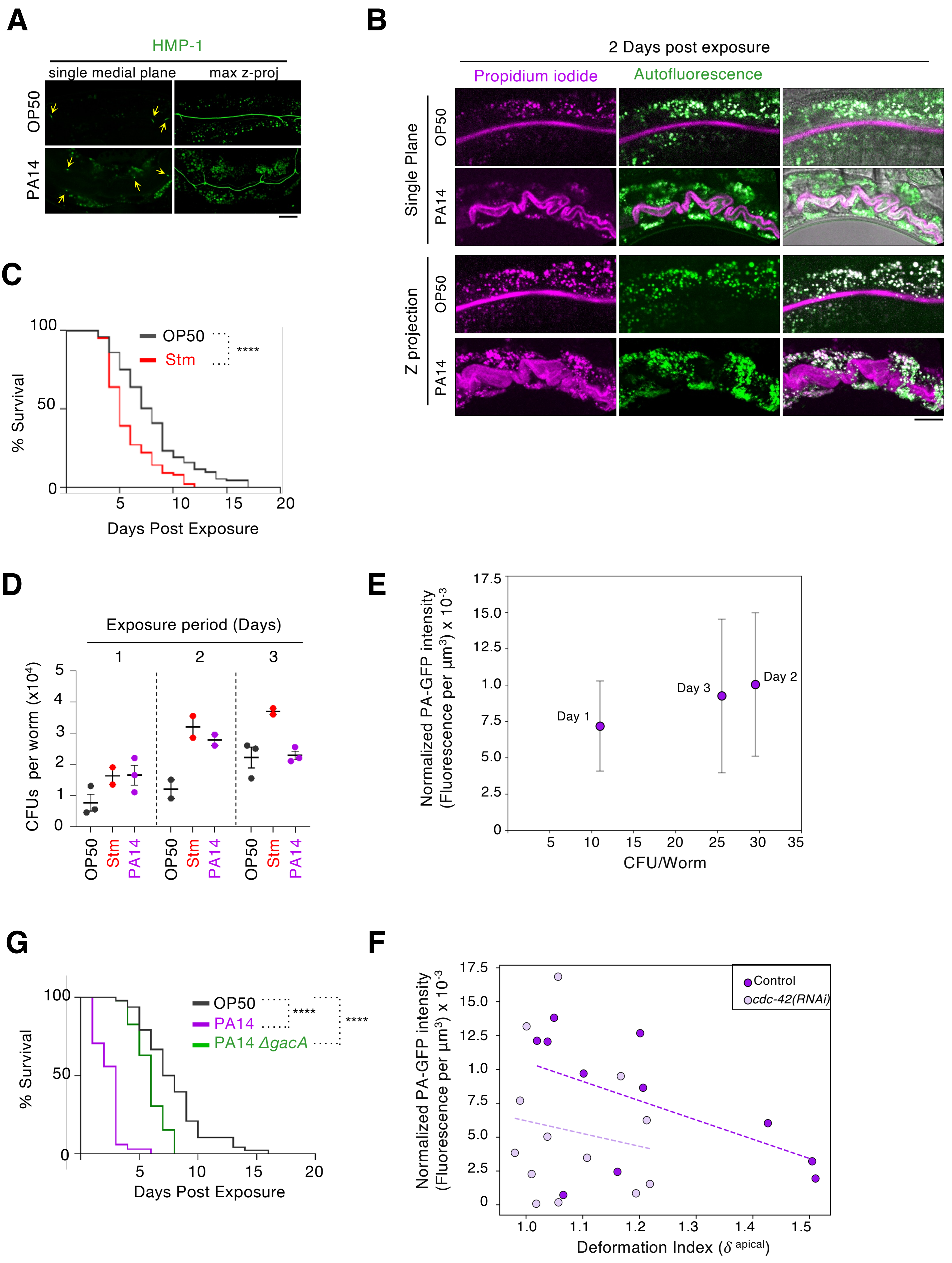
